# Nanoparticle-mediated delivery of PD-L1 inhibitor enhances γδ T cell immunotherapy against metastatic ovarian cancer cells

**DOI:** 10.64898/2026.06.04.730154

**Authors:** Laura O’Conner, Jessica Eakins, Mark Bates, Ola Ibrahim, Cara Martin, Victoria Malone, Steven G. Gray, Feras Abu Saadeh, Hassan Rajab, Doug A. Brooks, Ciaran M. Maguire, Stavros Selemidis, Julie David, Elena Matsa, Sharon O’Toole, John J. O’Leary, Derek G. Doherty, Bashir M. Mohamed

## Abstract

**Introduction:** Immune checkpoint inhibitors (ICIs) have only shown limited efficacy for patients with ovarian cancer (OC), partly due to the immune suppressive tumour microenvironment (TME) and platelet cloaking of the cancer cells. We tested a nanomedicine strategy to enhance γδ T cell immunotherapy by conjugating the PD-L1 inhibitor (BMS202) to nanodiamonds (NDs).

**Methods:** Patient-derived ascites cells were exposed to activated platelets to model platelet cloaking and the immunosuppressive phenotype in metastatic OC. ND/BMS202 nanocomplexes were then applied to platelet-conditioned OC cells and co-cultured with expanded γδ T cells. Cytotoxicity and immune activation were assessed by quantifying Granzyme B, CD107a, γ-H2AX, cleaved caspase-3, and cleaved caspase-8.

**Results:** ND-mediated delivery of BMS202 significantly enhanced γδ T cell-mediated killing of platelet cloaked OC cells in a dose-dependent manner, with greater efficacy than free BMS202 This enhanced cytotoxicity was supported by increased degranulation (CD107a), Granzyme B release, tumour cell apoptosis (caspase cleavage) and DNA damage (γ-H2AX staining).

**Conclusions:** ND-based delivery of the PD-L1 inhibitor (BMS202) enhances γδ T cell-mediated killing of platelet-cloaked metastatic OC cells. While our data support enhanced γδ T cell cytotoxicity following BMS202 delivery, direct evidence of PD-L1 target engagement or PD-1/PD-L1 binding inhibition was not demonstrated in this study. These findings nonetheless justify further validation in patient-derived organoid models to optimize this γδ T cell-based combination immunotherapy and advance its development as a precision therapeutic strategy for metastatic OC.

## Introduction

Ovarian cancer (OC) remains one of the most lethal gynaecological malignancies, largely because most patients present with advanced-stage metastatic disease. (1,2). Although prognosis is more favourable when OC is detected early, most patients present with advanced or metastatic disease, where the five-year survival rate is only around 44% (3,4). Metastatic OC initially spreads predominantly within the peritoneal cavity, however, cancer cells can also intravasate, enter the bloodstream and travel to distant sites where they can extravasate and lodge in other tissues, such as the lipid rich omentum within the abdominal cavity (5–8). Following intravasation, circulating metastatic OC cells can interact with several cell types, including cancer-associated fibroblasts, immune cells, endothelial cells and platelets which can support the transport and survival of these cancer cells within the bloodstream (9).

Platelets are particularly effective at shielding circulating tumour cells (CTCs), a phenomenon known as platelet cloaking. Platelet cloaking can protect CTCs from the immune system, promote epithelial-mesenchymal transition (EMT), and help them withstand circulatory shear forces (10). Furthermore, platelet cloaking can increase PD-L1 expression on CTCs, which can then interact with PD-1 on activated T-cells to suppress T-cell proliferation, cytokine production and cancer cell cytotoxicity (12). Platelets can also recruit granulocytes, including neutrophils, to further promote metastatic potential, drive angiogenesis and augment the immunosuppressive environment (11).

Because metastasis and chemotherapy resistance are so common in patients with OC, the disease remains difficult to treat (14). Patient survival has improved with the implementation of maintenance therapies, such as VEGF-A inhibitors and PARP inhibitors for patients with *BRCA* mutations or homologous recombination deficiency (15–17). Despite substantial efforts to develop immunotherapies for patients with OC, immune checkpoint inhibitors are yet to be approved for routine clinical practice. Pembrolizumab did show limited efficacy against recurrent or metastatic OC in the KEYNOTE-100 trial (18,19) and cell-based immunotherapies have shown encouraging safety and tolerability profiles in early clinical studies (20).

Activated immune cells, including γδ T cells, express cell surface PD-1 to control excessive immune activation and protect healthy tissue (22,21) from immune-mediated damage. Tumour cells exploit this biology by upregulating the expression of PD-L1 on their cell surface, which binds T-cell PD-1 to induce inhibitory signalling (23). This reduces γδ T cell IFN-γ production and cytotoxic activity (24,25). While the presence of intratumoral γδ T cells usually correlates with a more favourable prognosis (26), high PD□L1 expression on the tumour cells can negate this by inhibiting the activation and effector function of these T-cells (27,28).

Nanomaterials, including nanodiamonds (NDs), are being developed for cancer therapy to improve drug solubility, reduce systemic toxicity, enhance tolerability, and support localised drug delivery (29,30). For example, NDs have been modified to deliver doxorubicin to cancer cells, lowering the dose required and limiting associated side effects (31). In a previous study, ND-BMS202 conjugates were found to significantly augment immune□cell activity against malignant melanoma cells (32). BMS202 functions by binding to PD□L1 and promoting its dimerisation, which occludes the hydrophobic PD□1 interaction site (33).

This provides a strong mechanistic rationale for using NDs to concentrate BMS202 at the tumour–immune cell interface, where platelet cloaking and tumour-cell PD-L1 expression may otherwise limit γδ T-cell engagement and cytotoxicity. Here, this ND-based delivery platform has been adapted to develop a potential metastatic ovarian cancer therapeutic. The key point is that NDs help concentrate BMS202 precisely at the tumour–immune cell interface — the critical site where platelet cloaking physically obstructs γδ T-cell access and where tumour-cell PD-L1 expression actively suppresses their cytotoxic activity. This gives the nanodiamond strategy a clearer mechanistic rationale beyond general drug-delivery benefits, directly addressing the dual barriers of physical occlusion and immune checkpoint-mediated inhibition in the platelet-cloaked OC microenvironment.

## Materials and Methods

### Ethics statement

The Research Ethics Board at St. James’s Hospital and Adelaide and Meath Hospitals (SJH/AMNCH) approved this study (2012/11/04). Anonymised healthy female donor buffy coats were obtained from the Irish Blood Transfusion Service (IBTS, St. James’s Hospital), and PBMC were isolated by standard density gradient centrifugation. All experiments followed the Helsinki Declaration and the relevant institutional guidelines.

### Ovarian cancer cell isolation and culture

OC cells were isolated from deidentified patient ascites samples and designated as OCAS7 and OCAS8. Cells were cultured in RPMI 1640 (GIBCO, Invitrogen, Ireland) containing 10% (v/v) foetal calf serum, 20□mM HEPES, 10□μM nicotinamide, 10□μM SB202190, 1.25□mM N-acetyl-L-cysteine, 10□ng/mL FGF-10, 1□ng/mL FGF-2, 1× B27 supplement, 1:100 (v/v) Primocin, 10□μM Y-27632, 2□mM L-glutamine and 100□U/mL penicillin-streptomycin (Invitrogen, Ireland) at 37□°C in 5% CO□. After four days, unattached cells were removed by washing with phosphate buffer saline (PBS) and fresh medium was added (34,35). Cells were used for downstream experiments after a minimum of ten passages.

### Immunophenotypic profiling of OC cells

Patient-derived ascites cells (OCAS7 and OCAS8) were seeded in 96-well culture plates for 24□h, fixed with 3% paraformaldehyde (PFA) and washed with PBS (Sigma-Aldrich, Ireland) prior to staining with mouse monoclonal anti-p16, anti-WT1, anti-PAX8, anti-PD-L1, (ThermoFisher Scientific, Dublin, Ireland) and anti-β-Catenin, anti-thrombin, and anti-N-cadherin antibodies (Santa Cruz Biotechnology Inc., Germany). Samples were subsequently incubated with a goat anti-mouse FITC secondary antibody and nuclei were counter stained with Hoechst 33342 (1:1000). An EVOS™ M5000 Imaging System (ThermoFisher Scientific, Massachusetts, USA) was used for image acquisition.

### Nanocomplex development and characterisation

BMS202 (Cat. No. S7912-SEL, 10 mM in 1 mL DMSO) was purchased from Stratech (Ely, UK). ND powder (100 μg; gift from Nanodiamond Products Limited, Ireland, now Hyperion Materials & Technologies) was resuspended in deionized water (DW) to form an ND solution, and the suspension was autoclaved. Purified NDs were PEGylated using 0.2 mM 2-methoxy(polyethyleneoxy)propyl trimethoxysilane (Cat. No. S2535, Cymit Química S.L., Barcelona, Spain). As previously described (32), PEGylated NDs were mixed with 50 mM 2-(N-morpholino)ethanesulfonic acid (MES) buffer (1.0 M, Cat. No. J60763.AK, ThermoFisher) pH 6, for 24 h. The ND-PEG particles were then concentrated and resuspended in 0.5 mL MES buffer containing 10 mg of N-ethyl-N’-[3-dimethylaminopropyl]carbodiimide (EDC, Cat. No. PG82079, ThermoFisher Scientific) and 10 mg of N-hydroxysuccinimide (NHS, Cat. No. 24500, ThermoFisher Scientific), followed by vigorous agitation for 15 min. The reaction mixture was washed twice by centrifugation at 13,300 rpm (16,100 x g) with DW, then resuspended in 1 mL MES buffer containing 100 μL of 10 mM BMS202. The mixture was agitated for 4□h at room temperature. Particles were washed again with DW and resuspended in purified DW to yield the nanocarrier complex, designated ND/BMS202 **(Fig.□1)**. ND and ND-PEG-BMS202 were physicochemically characterised using a NanoSight Pro system (Malvern Panalytical Ltd, Malvern, United Kingdom) for particle size and concentration (10-1000□nm range).

### Platelet preparation

Blood was collected into 10% ACD and rocked at 30□rpm for 15□min. Platelet-rich plasma (PRP) was obtained by centrifugation at 170×g for 10□min at room temperature. The PRP was acidified to pH□6.5 with ACD, and prostaglandin E2 (1□μM final) was added. Platelets were then pelleted at 720×g for 10□min, resuspended in JNL buffer (130□mM NaCl, 10□mM sodium citrate, 9□mM NaHCO□, 6□mM D-glucose, 0.9□mM MgCl□, 0.81□mM KH□PO□, 10□mM Tris, pH□7.4), and counted on a Sysmex XP-300 haematology analyser (Sysmex Corporation, Kobe, Japan). Before use, platelets were activated with 0.18□mM CaCl□. Activated platelets were added to cell samples within 60□minutes of blood collection.

### Assessment of EMT in OC cells after platelet co-culture

To induce a more metastatic phenotype, 5,000 OCAS7 or OCAS8 cells were seeded per well in 96-well culture plates and exposed to activated platelets at a ratio of 2000 platelets per cancer cell (2000:1 platelet:cancer cell ratio) for 48□h. EMT status was evaluated by immunohistochemistry, and platelet binding was verified using an anti-GPIbα antibody (Clone HIP1, BD Biosciences).

### **γδ** T cell isolation, expansion and maintenance

PBMCs were isolated from buffy coats by density gradient centrifugation over Lymphoprep. γδ T cells were enriched from 2×10□ PBMCs by negative selection using the CliniMACS TCRα/β Biotin Kit (Miltenyi Biotec). Cells were Fc blocked, labelled with biotinylated anti TCRα/β antibody, and depleted using anti biotin magnetic beads. The αβ depleted fraction (containing γδ T cells, B cells, NK cells, and monocytes) was then stimulated on anti CD3 (OKT3)–coated plates in the presence of IL 2 and IL 15. This culture system drives robust expansion of Vδ1, Vδ2, and Vδ3 subsets; non proliferating cells are lost within one week, yielding tens of millions of highly enriched γδ T cells within 2–3 weeks. Because γδ T cells are not MHC restricted, they were used with ovarian cancer samples from allogeneic donors. Previous studies have confirmed that expanded γδ T cells lack significant alloreactivity against allogeneic tumour cells due to their MHC-independent recognition mechanisms, supporting their use across donor–recipient mismatches (20).

### Immunophenotyping of **γδ** T cells by flow cytometry

Expanded γδ T cells were stained with a viability dye (eFluor 506 Fixable Viability Dye), followed by an Fc blocking agent and antibodies against CD3, Vδ1, Vδ2, Vδ3 and PD-1. Samples were run on a BD FACSCanto II or LSR Fortessa and analysed using FlowJo software to get subset frequencies.

### Experimental co-culture design

After platelet induced EMT, OCAS7 and OCAS8 cells were seeded overnight (5,000 cells per well in 96 well culture plates or 100,000 cells per well in 6 well culture plates). The following day, cells were treated with either free BMS202 or ND/BMS202 complexes for 6**□**h, after which γδ T cells were added at a 10:1 effector to target ratio and co cultured overnight **(Fig. 1)**.

**Figure 1.**
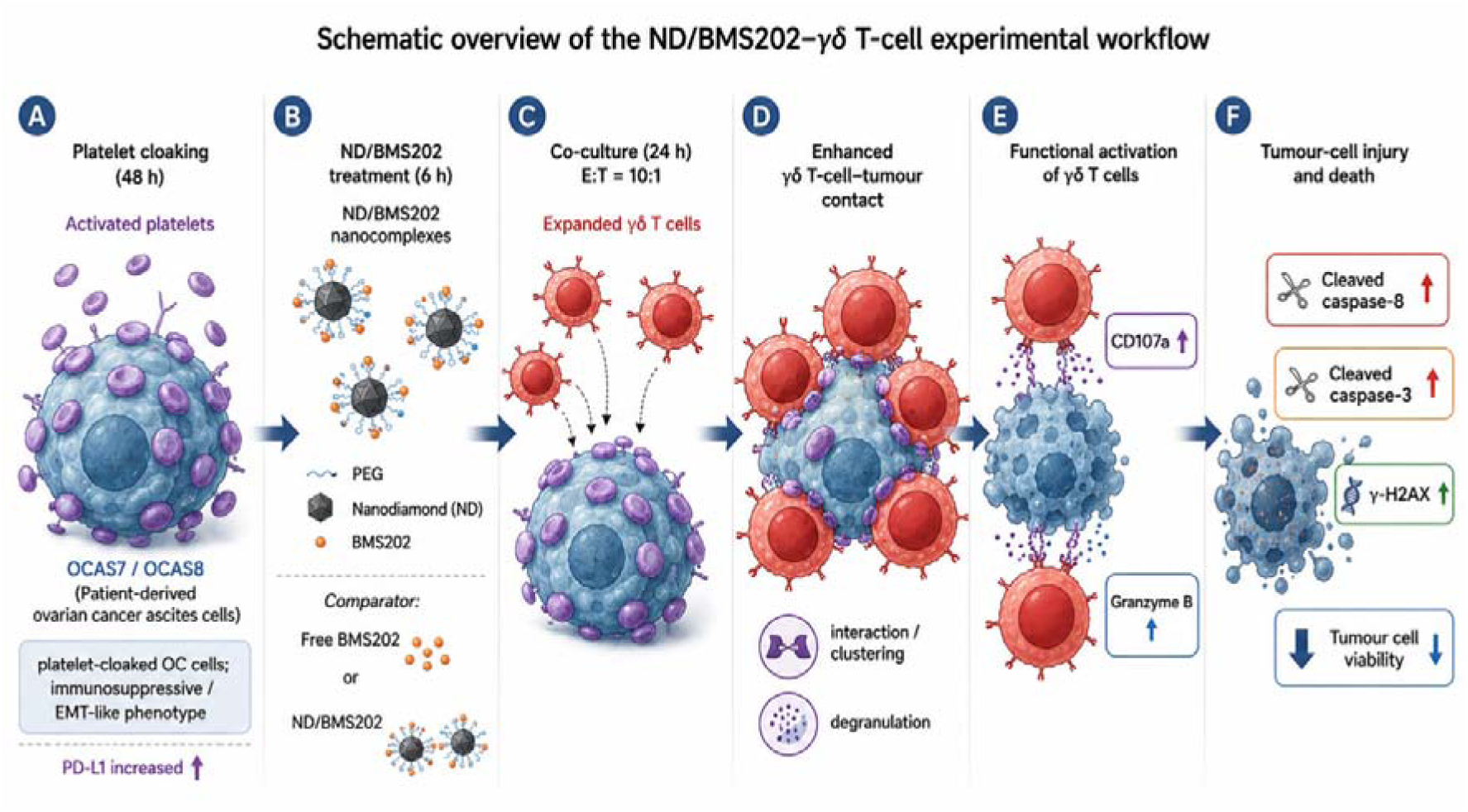
Schematic overview of the ND/BMS202–γδ T-cell experimental workflow. Patient-derived OC ascites cells (OCAS7/OCAS8) were exposed to activated platelets for 48 h to generate platelet-cloaked OC cells with an immunosuppressive, EMT-like phenotype. Cells were then treated for 6 h with ND/BMS202 nanocomplexes, or comparator treatments, before co-culture with expanded γδ T cells at a 10:1 effector-to-target ratio for 24 h. ND-mediated delivery of BMS202 enhanced γδ T-cell–tumour cell contact and promoted γδ T-cell functional activation, as shown by increased CD107a expression and Granzyme B release. This was associated with increased tumour-cell injury and death, including elevated cleaved caspase-8, cleaved caspase-3 and γ-H2AX, together with reduced tumour-cell viability.

### Assessment of **γδ** T cell–tumour cell interactions

OCAS7 and OCAS8 were treated with 10**□**μM free BMS202 or ND/BMS202 (10**□**μM BMS202 equivalent) for 6**□**h, then added γδ T cells (10:1) overnight. Plates were gently washed to keep non-adherent T cells, fixed with 3% PFA, and stained with FITC-conjugated anti-CD45 (ThermoFisher Scientific) for γδ T cells and TRITC-labelled phalloidin (Sigma-Aldrich, Ireland) for OC cells. T cells that contacted OC cells and solitary T cells were counted using an EVOS™ M5000 Imaging System (ThermoFisher Scientific).

### Analysis of Granzyme B and CD107a expression

After co-culture γδ T cells were harvested. Granzyme B (GrB) was measured by high-content imaging on a Cytell™ system (Cytely AB, Lund, Sweden). CD107a was measured by flow cytometry after staining with anti-CD107a antibody, which was added at the time the γδ T cells were added to the OC cells.

### Cytotoxicity analysis

Cytell system was used in this study to quantify cytotoxicity. Cells were fixed with 3% PFA after treatment and co-culture, then stained with antibodies against cleaved caspase-3, cleaved caspase-8 and γ-H2AX. Fluorescence intensity per well was analysed by high-content imaging on a Cytell™ system (Cytely AB).

### Statistical analysis

GraphPad Prism 8 was used for all statistical analyses. Data are presented as mean ± SEM from three independent biological replicates (n=3 per condition). Normality and homogeneity of variances were assessed where applicable; parametric tests were used unless assumptions were violated. Comparing multiple treatments against a single control (e.g., all treatments vs. NT control), one-way ANOVA followed by Dunnett’s post hoc test was used. For experiments requiring comparisons between all pairs of treatments (e.g., ND/BMS202 doses vs. free BMS202 vs. ND alone), one-way ANOVA with Tukey’s post hoc test was applied. Pairwise comparisons between two groups were performed using an unpaired two-tailed Student’s t test. A p value < 0.05 was considered statistically significant (*p < 0.05, **p < 0.01, ***p < 0.001). Specific comparisons for each figure are detailed in the figure legends and supplementary tables.

## Results

### Characterisation of OC cells after isolation and expansion

Patient-derived ascites cells (OCAS7 and OCAS8) were phenotypically characterised using immunohistochemical markers standard for OC diagnosis in the Histopathology Laboratory (St. James’s Hospital). The panel included PAX8, a sensitive and specific marker for tumours of Müllerian origin (36); WT1, whose expression varies diagnostically among OC histological subtypes (37); and p16, a tumour suppressor protein overexpressed in subtypes such as high-grade serous OC (38). Both OCAS7 and OCAS8 demonstrated positive staining for PAX8, WT1 and p16, supporting their identity as OC cells (Fig. 2a). PD-L1 expression was also detected in both cell lines (Fig. 2b).

**Figure 2 (a).**
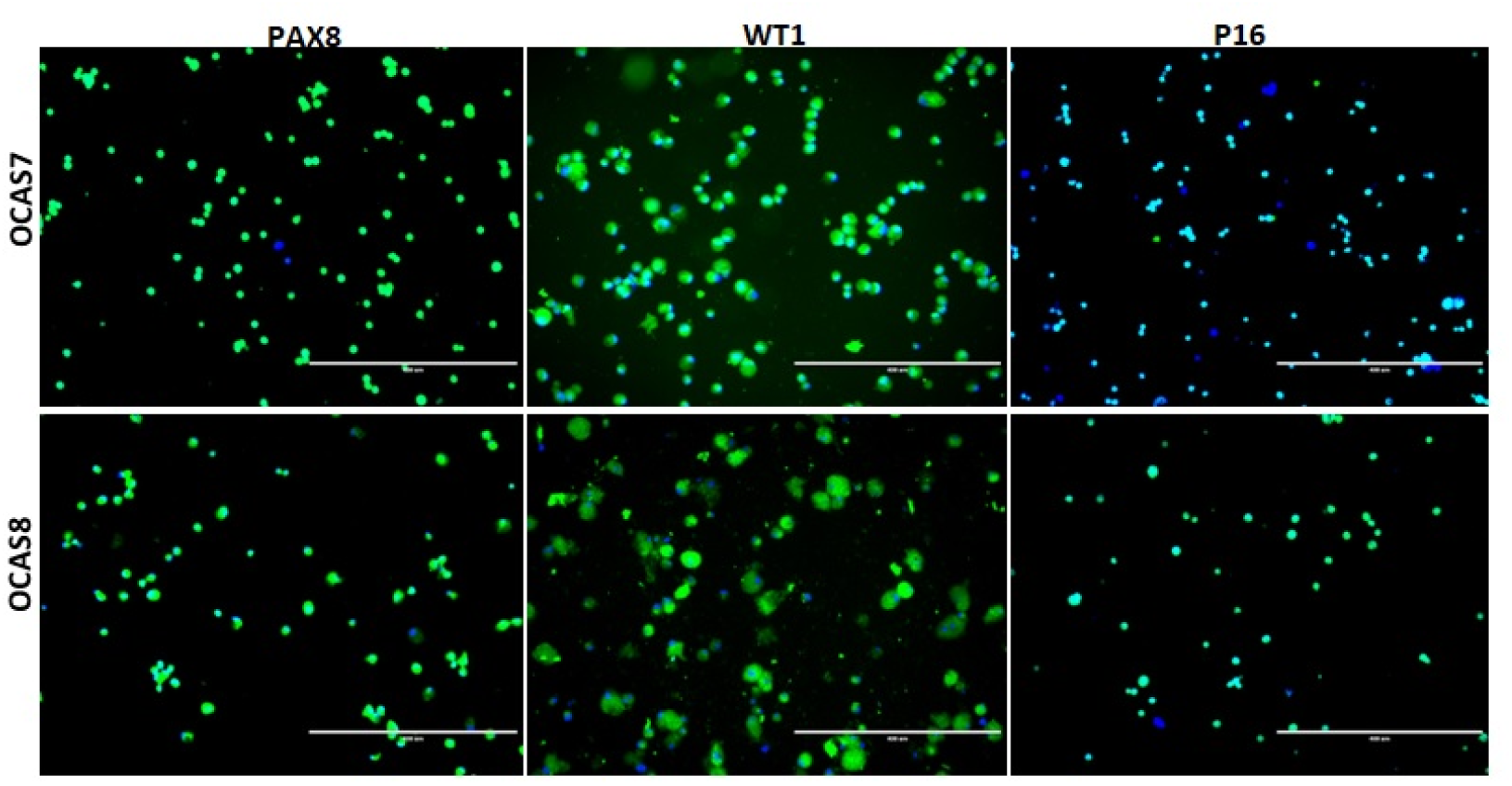
Representative image of OCAS7 and OCAS8 cells. OC cells were isolated and expanded from OC patients’ ascites. Cells were labelled with anti-PAX8, anti-WT1 and anti-P16 antibodies, using FITC labelled secondary antibody (green) and cells counterstained with Hoechst 33342, (blue) for visualisation of the nuclei. Imaging was performed using an inverted microscope (10X), and scale bar represents 400 µm.

**Figure 2 (b).**
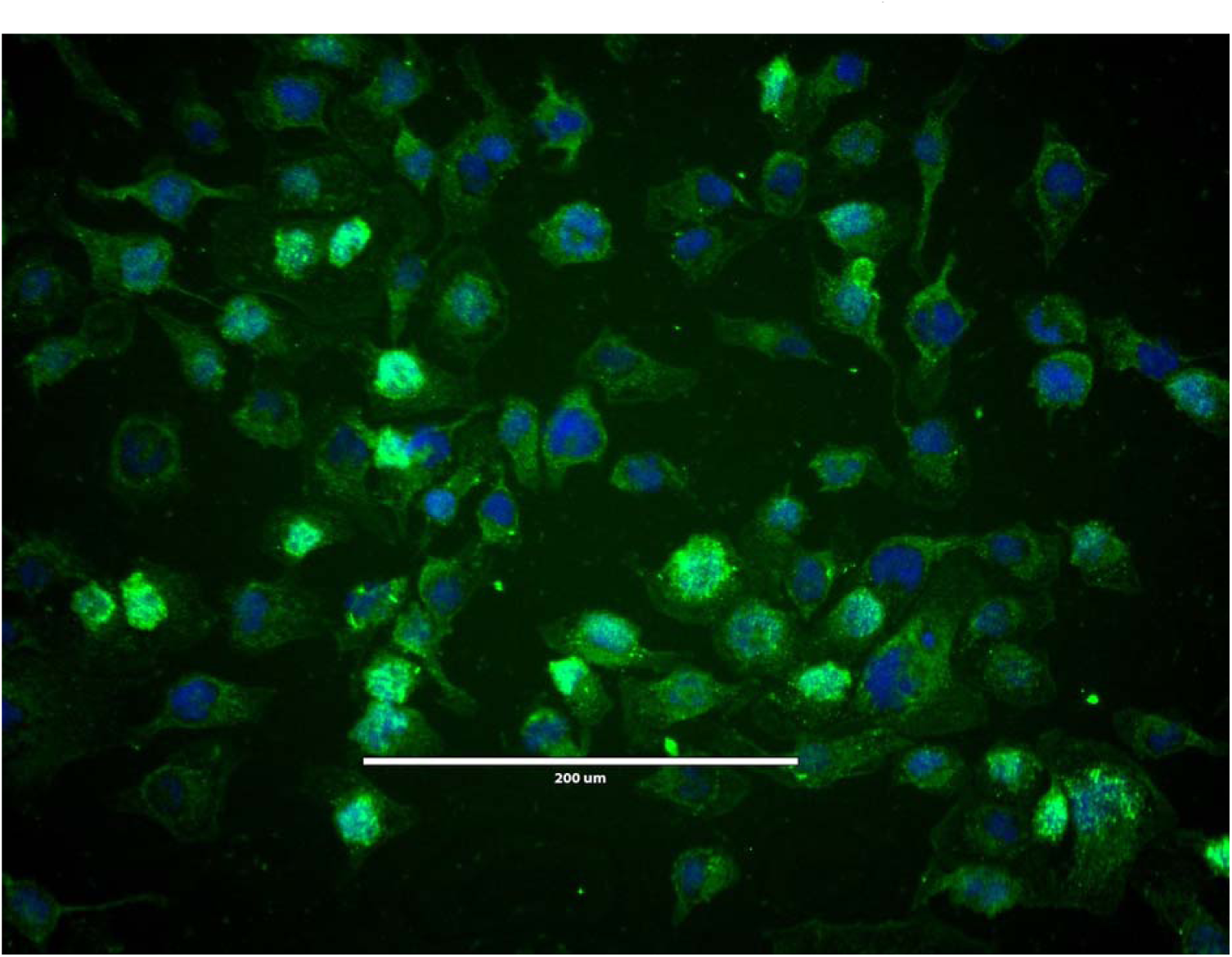
Representative image of PD-L1 expression. OC cells were isolated and expanded from OC patients’ ascites. Cells labelled with anti-PD-L1 anti-antibodies and FITC ladled secondary antibody (green), and cells counterstained with Hoechst 33342, (blue) for visualisation of the nuclei. Imaging was performed using an inverted microscope (20X), and scale bar represents 200 µm.

### Platelet-induced cloaking and EMT phenotype induction in OC cells

After 48**□**h co-culture with platelets at a ratio of 2000 platelets per cancer cell (2000:1 platelet:cancer cell ratio), the OC cells were heavily cloaked with platelets, as indicated by strong anti-CD42b (GPIbα) staining (PE). Concurrent HER2 staining (FITC) confirmed the identity of the OC cells (Fig.□3). Platelet exposure also induced a more mesenchymal phenotype, characterised by increased β-catenin, thrombin, and N-cadherin expression, reduced E-cadherin expression, and altered cell morphology (Fig.□4).

**Figure 3.**
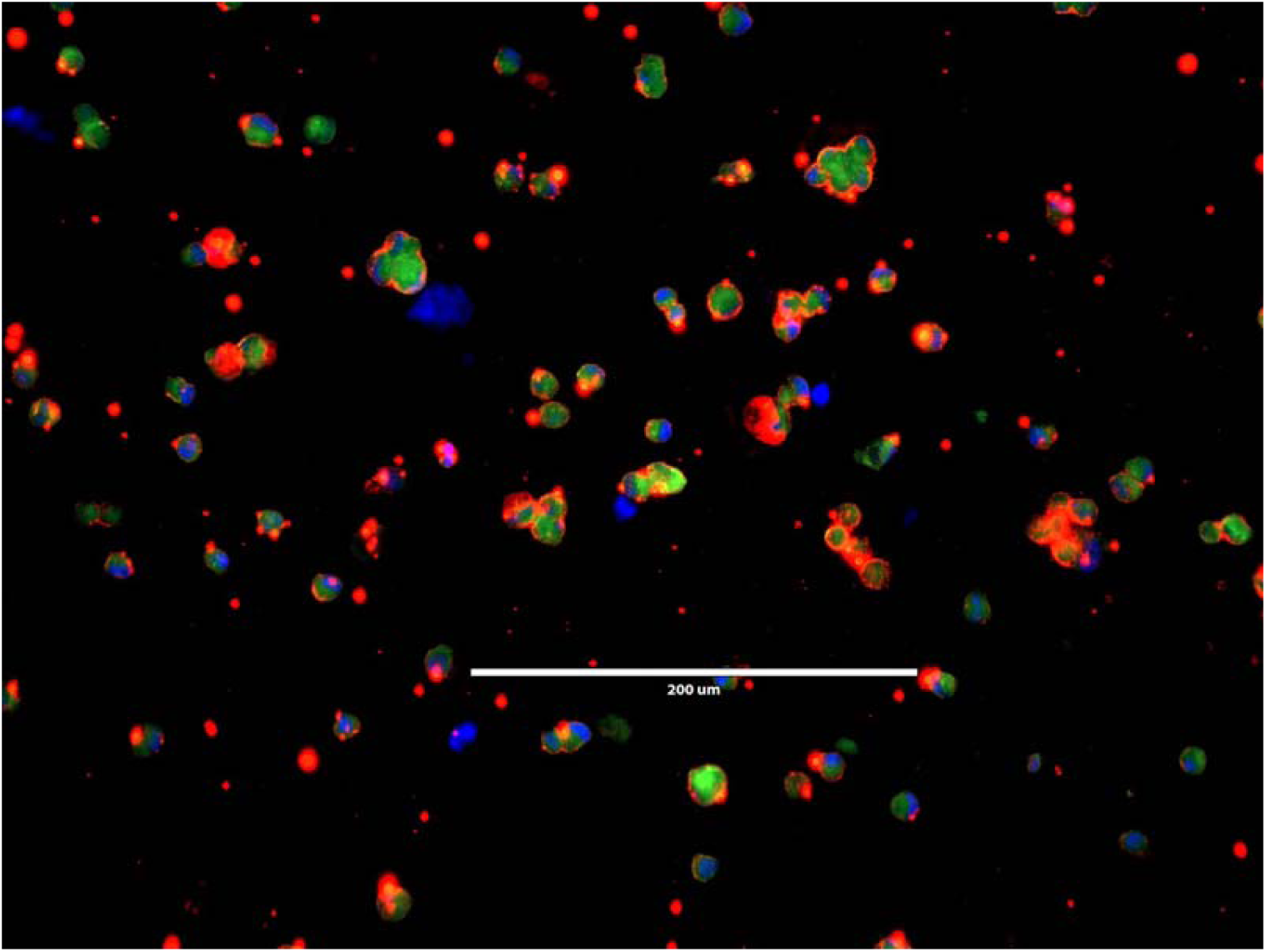
Representative image of OC cells cloaked with platelets. OC cells were labelled with anti-CD42b antibodies and FITC conjugated-HER2 antibodies for 24 h. Cells then were then labelled with TRITC-conjugated secondary antibodies antibody (red) and cells counterstained with Hoechst 33342, (blue) for visualisation of the nuclei. Imaging was performed using an inverted microscope (10X), and scale bar represents 400 µm.

**Figure 4a.**
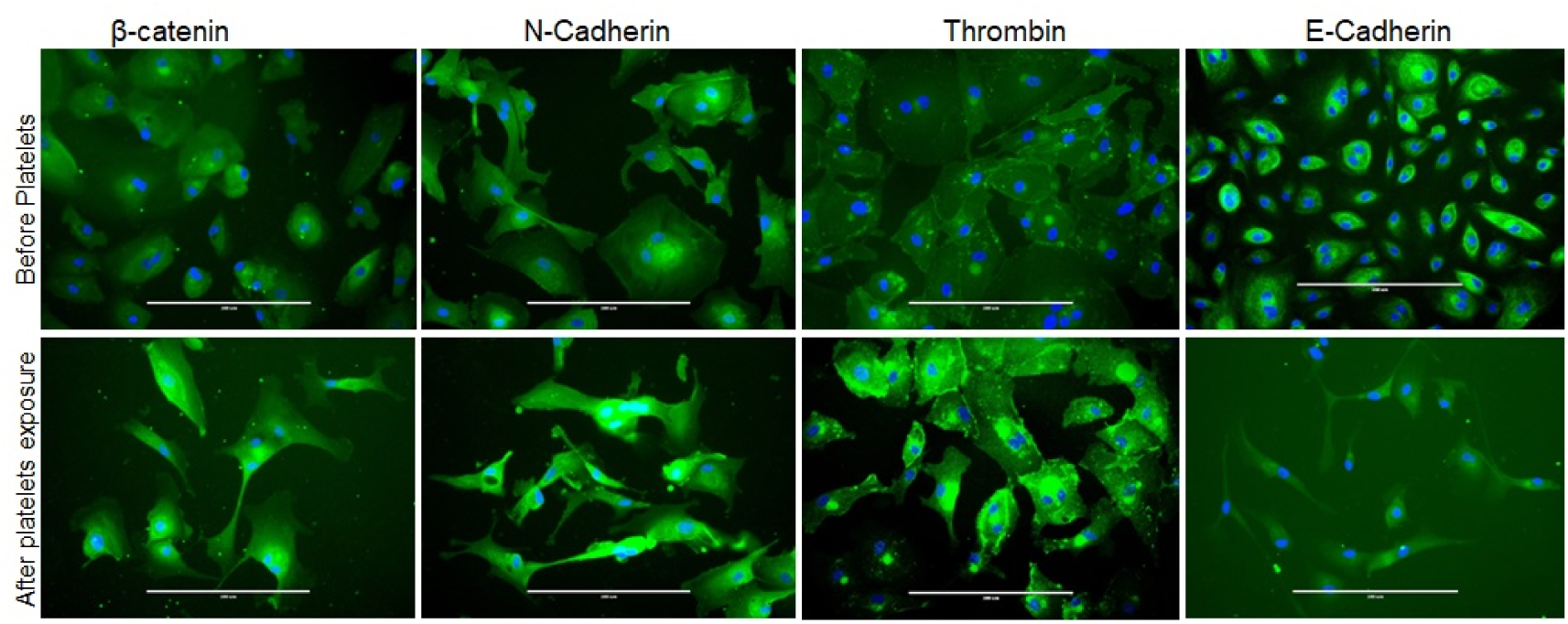
Representative image of OC cells pre and post platelets cloaking. OC cells were labelled with anti-β-catenin, N-Cadherin, Thrombin and E-Cadherin antibodies for 24 hand FITC-conjugated secondary antibodies (green), and cells counterstained with Hoechst 33342, (blue) for visualisation of the nuclei. Imaging was performed using an inverted microscope

### Flow cytometric profiling of **γδ** T cell subsets

Following isolation from PBMCs, γδ T cells were stained with fluorescently conjugated antibodies against CD3 (Pacific Blue), Vδ3 (APC), Vδ1 (APC/Vio770), and Vδ2 (FITC). Flow cytometric analysis was performed after gating on lymphocytes and excluding doublets and dead cells. The expanded γδ T-cell population contained Vδ1**□**, Vδ2**□** and Vδ3**□** subsets, representing 24.0%, 14.3% and 9.5% of total CD3**□** cells, respectively (Fig.**□**5a-c).

**Figure□5.**
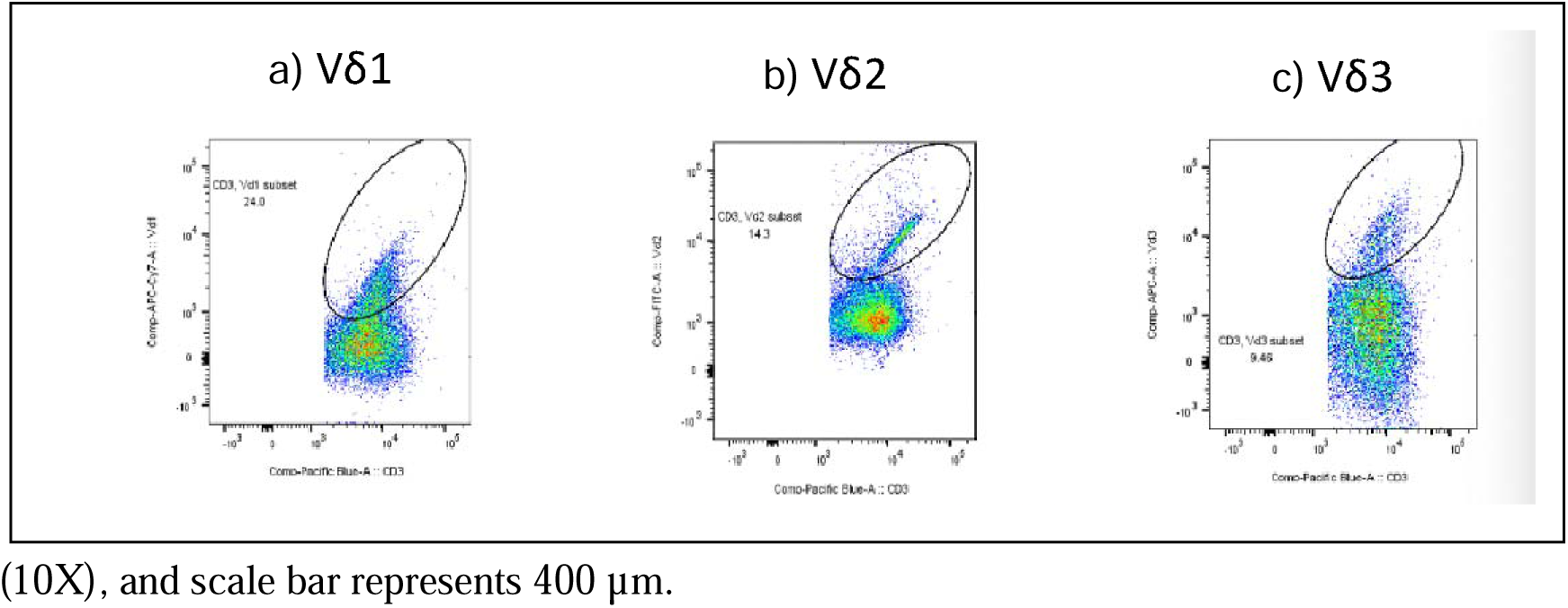
γδ T cell subset analysis by flow cytometry. (a) Gating strategy for CD3**□** lymphocytes. (b) Representative plots showing Vδ1, Vδ2, and Vδ3 expression. (c) Subset frequencies as percentage of CD3**□** cells (mean ± SEM, n=3). Vδ1**□**: 24.0%, Vδ2**□**: 14.3%, Vδ3**□**: 9.46%.

### Hydrodynamic Size Characterisation of ND-PEG-BMS202 Nanocomplexes

The hydrodynamic diameter of the fabricated nanocomplexes was measured in deionised water using nanoparticle tracking analysis on the NanoSight Pro system. Bare NDs displayed an average hydrodynamic diameter of 72**□**nm (Fig.**□**6a). Following PEGylation and conjugation with BMS202, ND-PEG-BMS202 nanocomplexes showed a concentration-dependent increase in mean particle size: 193**□**nm at 2.5**□**μM BMS202, 206**□**nm at 5**□**μM, and 222**□**nm at 10**□**μM. These data indicate successful formation of ND/BMS202 nanocomplexes with dose-dependent changes in particle size.

**Figure 6.**
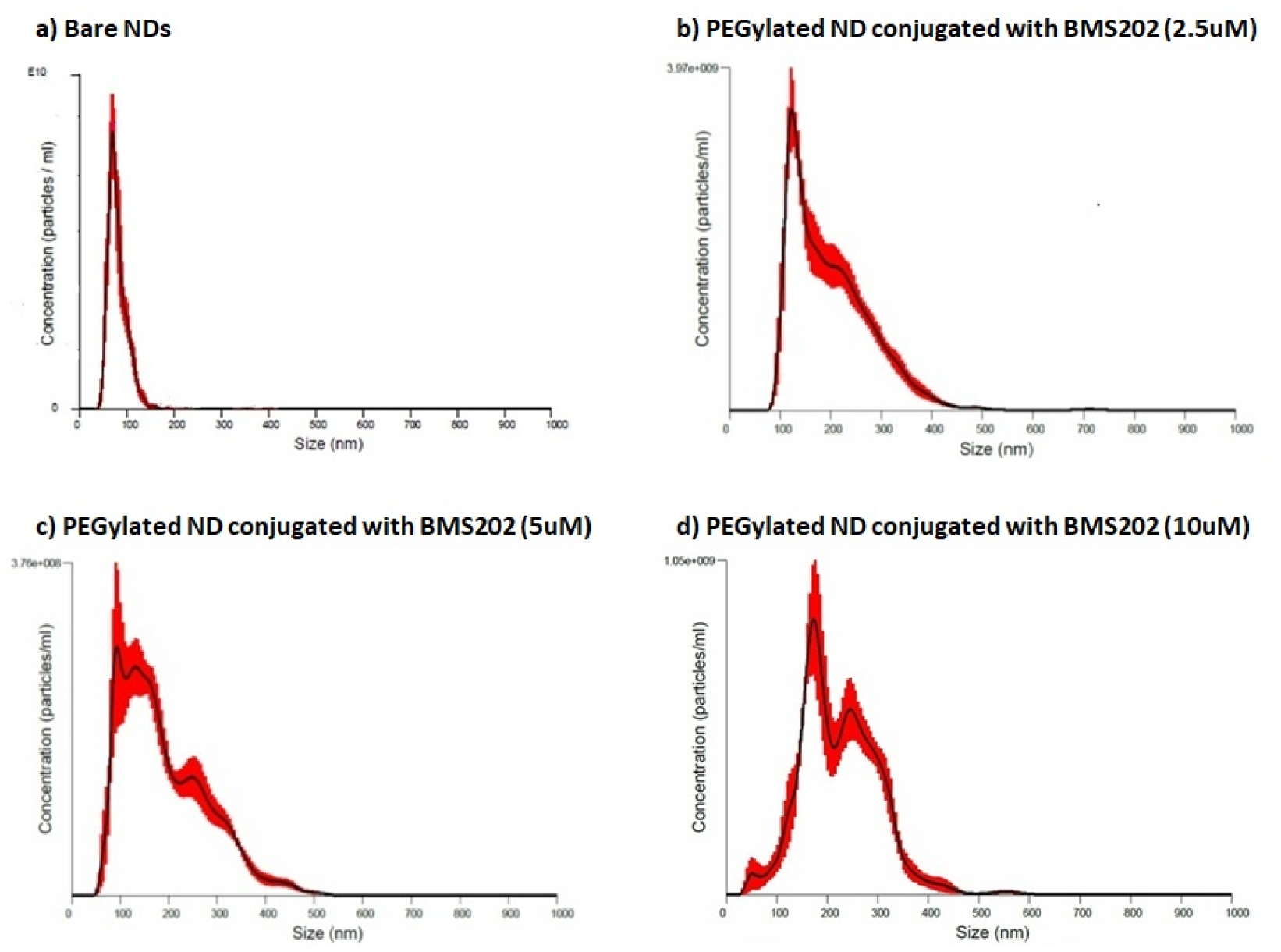
Nanoparticle tracking analysis (NTA) of size distribution. (a) Bare ND showing an average hydrodynamic diameter of 72 nm. (b–d) PEGylated ND conjugated with BMS202 resulting ND/BMS202 at increasing concentrations: (b) 2.5 μM PD-L1i – 193 nm; (c) 5 μM PD-L1i – 206 nm; (d) 10 μM PD-L1i –222 nm. Measurements were performed in deionized water. Data demonstrate a BMS202 (PD-L1i) concentration-dependent increase in the average size of ND/PD-L1i nanocomplexes compared to bare ND.

### ND/BMS202 enhanced **γδ** T cell–tumour interactions

OCAS7 and OCAS8 cells pre-treated with or without platelets were exposed to one of the following for 24 h: (i) our developed nanocomplex (ND/BMS202) at concentrations of 2.5 µM, 5 µM, or 10 µM (BMS202 equivalent), (ii) 10 µM BMS202 alone, or (iii) NT control. Cells were then co-cultured with γδ T cells at an effector-to-target (E:T) ratio of 10:1 for an additional 24 h. Following co-culture, cells were fixed with 3% PFA and stained. Cell nuclei were visualised with Hoechst 33342, while γδ T cells were specifically stained with an anti-CD45 antibody, and OC cells were stained with FITC-conjugated phalloidin to label F-actin (Fig. 7 a-f). For each treatment condition, a series of images was captured. Within each image, the number of γδ T cells in direct contact with OC cells and the number of solitary, non-interacting γδ T cells were counted.

**Figure 7.**
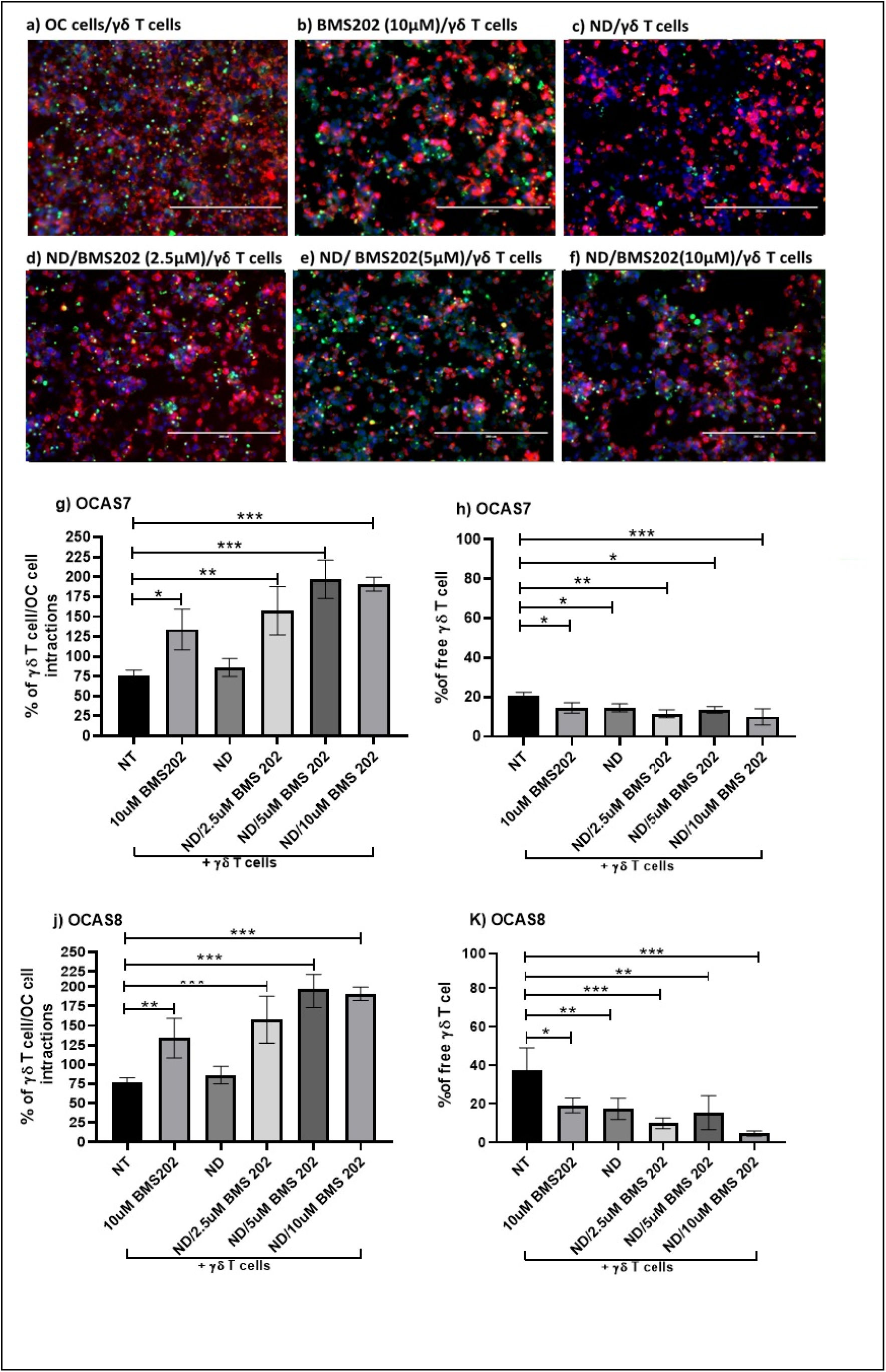
Representative images of γδ T cells and OC cell interactions. OC cells (**OCAS7 sample)** expanded from the tumour sample were exposed to platelets (at a ratio of 2000 platelets to 1 cancer cell) for 48h. a) OC cells were either co-cultured with γδ T cells (NT/γδ T cells) or (b), treated with10 μM BMS202 alone (c), ND (d), ND/2.5 μM BMS202 (e), ND/5 μM BMS202 and (f) ND/10 μM BMS202 for 6 h, and then co-incubated with γδ T cells (γδ T cells to 1 cancer cell) for an additional 24 h. Cells were washed in PBS and then fixed in 3% PFA, stained with FITC-conjugated anti-CD45 for γδ T cells and TRITC-labelled phalloidin for OC cells and counterstained with Hoechst 33342, (blue) for visualisation of cell nuclei. Imaging was performed using an inverted microscope (20x), and the scale bar represents 200 μm. Then the number of interacted (g&j) and free (h&k) γδ T cells were counted. Statistical significance was determined using one-way ANOVA followed by Dunnett’s post hoc test. The data were represented as mean ± SEM (n=3). “*” for p < 0.05, “**” for p < 0.01; and “***” for p < 0.001. vs. NT control.

ND/BMS202 treatment enhanced γδ T-cell interactions with platelet-cloaked OC cells. In OCAS7 cells, all tested concentrations of ND/BMS202 significantly increased the proportion of γδ T cells in direct contact with tumour cells compared with the NT control (2.5 µM: p<0.01; 5 µM: p<0.001; 10 µM: p<0.001; Fig. 7g). Free BMS202 also increased γδ T-cell interactions, although to a lesser extent (p<0.05). Similarly, in OCAS8 cells, ND/BMS202 significantly enhanced γδ T-cell interactions at all concentrations tested (all p<0.001), while free BMS202 also produced a significant increase (p<0.01; Fig. 7j). These effects were accompanied by a corresponding reduction in free, non-interacting γδ T cells across both cell lines.

### Granzyme B expression was increased by ND/BMS202 treatment

Granzyme B expression was significantly increased when γδ T cells were co-cultured with OC cells pre-treated with BMS202. This effect was observed in both patient-derived OC cell lines, OCAS7 and OCAS8, and was further enhanced when BMS202 was delivered using the ND nanocomplex.

For both OCAS7 and OCAS8, all BMS202-containing treatments significantly increased Granzyme B expression compared with the NT control. This included 10 µM free BMS202 and all tested concentrations of ND/BMS202. The response to ND/BMS202 was dose-dependent, with 2.5 µM producing a significant increase and the 5 µM and 10 µM doses producing stronger responses in both cell lines (Fig. 8g–h; Supplementary Tables 1 and 2). In contrast, naked NDs alone did not significantly alter Granzyme B expression, indicating that the effect was associated with BMS202 rather than the ND carrier.

**Figure 8.**
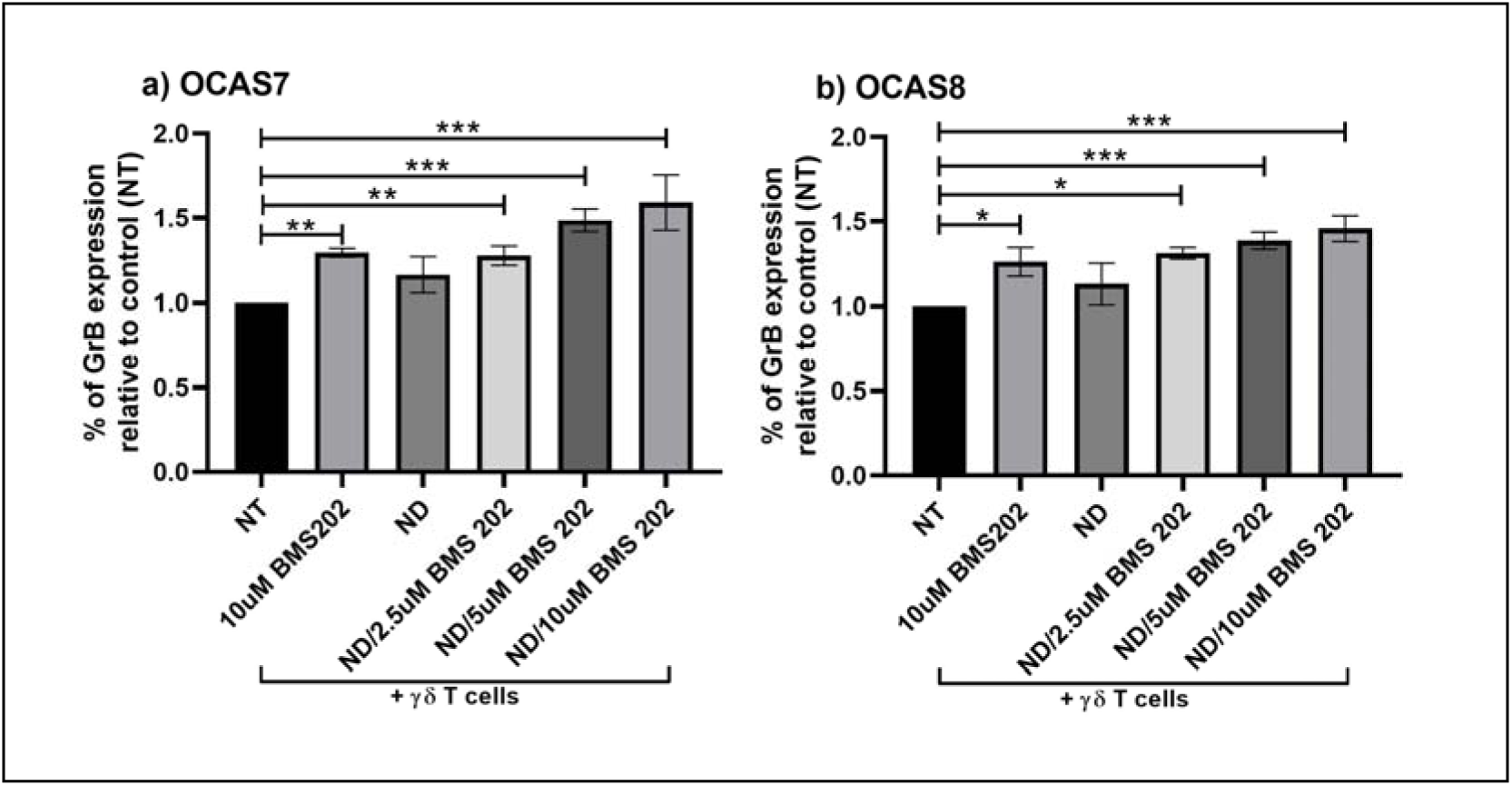
Detection of function al γδ T cell response to OC cells post-exposure to ND/BMS202. Quantification of Granzyme B for OCAS7 (a) and OCAS8 (b). Statistical significance was determined using one-way ANOVA followed by Dunnett’s post hoc test. The data were represented as mean ± SEM (n=3). “*” for p < 0.05, “**” for p < 0.01; and “***” for p < 0.001. vs. NT control.

The nanocomplex format demonstrated a distinct advantage, particularly in the OCAS8 model. For OCAS8 cells, the ND/10 µM BMS202 complex induced a significantly stronger GrB response than 10 µM BMS202 alone (p<0.05). Furthermore, for both cell lines, the higher-dose nanocomplexes (5 µM and 10 µM) triggered significantly greater GrB expression than the ND-only control (OCAS7: p<0.01 for both; OCAS8: p<0.01 and p<0.001, respectively).

In summary, these results demonstrate that the ND/BMS202 nanocomplex effectively and consistently enhanced the cytotoxic potential of γδ T cells against OC cells in a dose-responsive manner. The data suggest that the ND delivery platform can enhance the efficacy of the PD-L1 inhibitor, with this advantage being clearly evident in the OCAS8 cell line.

### CD107a degranulation was enhanced by ND/BMS202

CD107a expression, used as a marker of γδ T-cell degranulation, was significantly increased following co-culture with OC cells pre-treated with BMS202-containing formulations (Fig. 9a–f). This effect was observed in both OCAS7 and OCAS8 cells and was strongest when BMS202 was delivered using the ND nanocomplex. Pre-treatment with ND/BMS202 significantly increased γδ T-cell CD107a expression compared with the NT control at all tested concentrations in both cell lines (all p<0.001). Free BMS202 also significantly increased CD107a expression relative to the NT control (p<0.001). Naked NDs alone produced a small but significant increase in CD107a expression in the OCAS7 model (p<0.05), but this was not observed in OCAS8 cells. The highest tested dose of the ND/BMS202 nanocomplex (10 µM BMS202 equivalent) induced significantly greater CD107a expression than free BMS202 at 10 µM in both OCAS7 and OCAS8 cells (p<0.001). In OCAS7 cells, ND/BMS202 at 10 µM also induced significantly greater CD107a expression than ND/BMS202 at 2.5 µM or 5 µM (Fig. 9g–h; Supplementary Tables 3 and 4).

**Figure□9.**
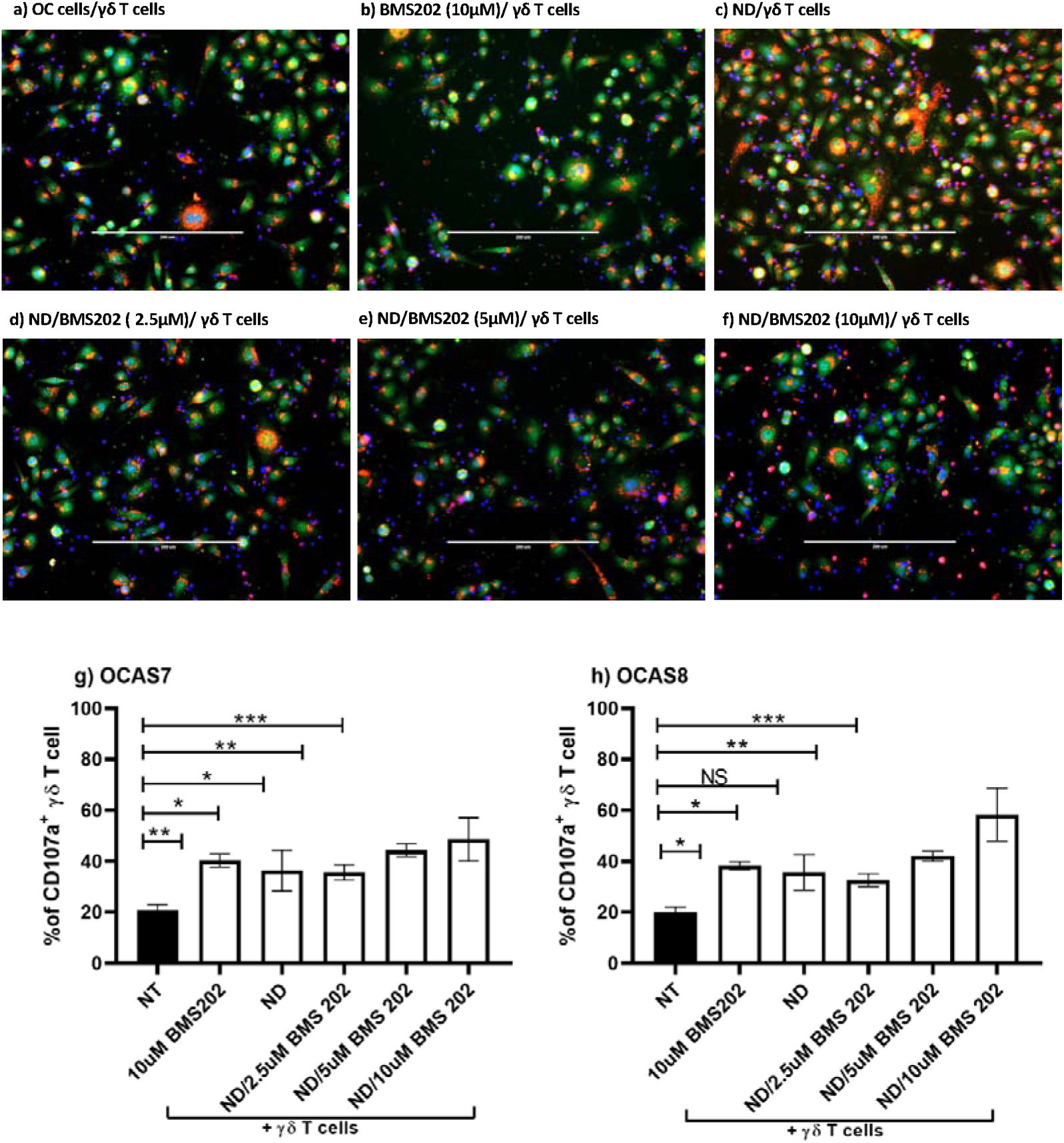
CD107a degranulation in γδ T cells. Representative images of CD107a (red) expression in γδ T cells co-cultured with OCAS8 cells. a) OC cells were either co-cultured with γδ T cells (NT/γδ T cells) or (b), treated with10 μM BMS202 alone (c), ND (d), ND/2.5 μM BMS202 (e), ND/5 μM BMS202 and (f) ND/10 μM BMS202 for 6 h, then cells were co-cultured with γδ T cells (10 γδ T cells to 1 cancer cell) for an additional 24 h. OC cells were probed with CD107a antibody (red) and counterstained using DNA Hoechst 33342 for visualisation of the nuclei (blue). Images were performed using an inverted microscope (20X), the scale bar represents 200 µm. Quantification of CD107a expression in OCAS7 (g) and OCAS8 (h). Statistical significance was determined using one-way ANOVA followed by Dunnett’s post hoc test. The data were represented as mean ± SEM (n=3). “*” for p < 0.05, “**” for p < 0.01; and “***” for p < 0.001. vs. NT control.

### ND/BMS202 enhanced **γδ** T-cell-mediated apoptosis, DNA damage and killing of OC cells

To determine how ND/BMS202 nanocomplexes modulate γδ T-cell cytotoxicity, the apoptotic activation (cleaved caspase-8 and caspase-3), DNA damage (γ-H2AX), and overall cell killing in OCAS7 and OCAS8 cells following treatment and γδ T-cell co-culture was measured. Across both derived cells, ND/BMS202 nanocomplexes markedly enhanced apoptotic signalling. Medium- and high-dose nanocomplexes (5**□**µM and 10**□**µM BMS202 equivalent) significantly increased cleaved caspase-3 expression relative to NT controls (p<0.001), with 10**□**µM free BMS202 producing a similar but less pronounced effect. The 2.5**□**µM nanocomplex dose selectively increased caspase-3 in OCAS8 cells (Fig. 10 a-h and Supplementary Table 5 & 6). All nanocomplex concentrations robustly activated cleaved caspase-8 (p<0.01-0.001), whereas naked NDs had no effect (Fig. 11 a-h and Supplementary Table 7 & 8). . No significant differences were observed between 10**□**µM free BMS202 and any nanocomplex dose for caspase-3 induction.

**Figure 10.**
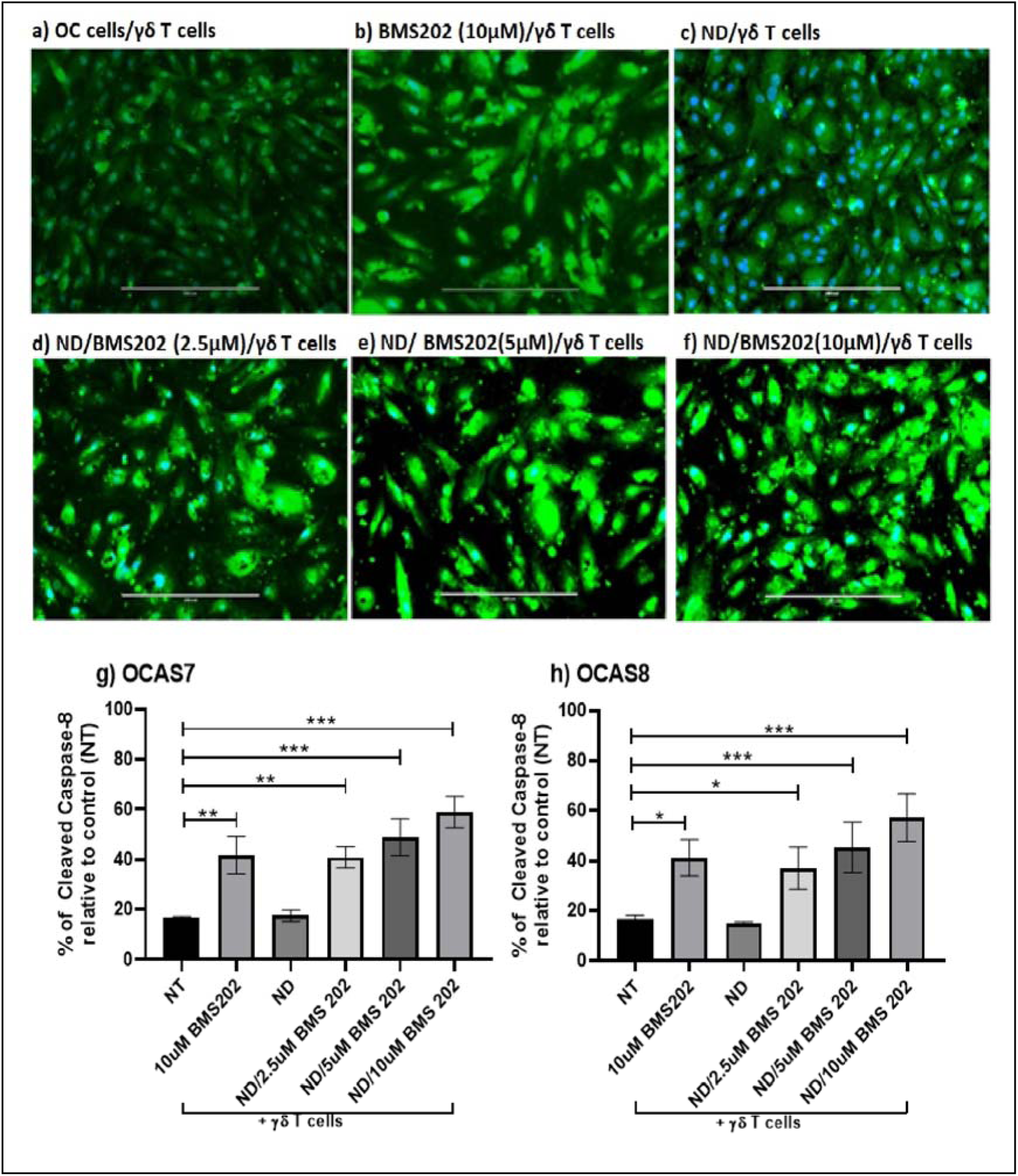
Cleaved caspase-3 expression in OC cells after ND/BMS202 and γδ T cell coculture. (a–f) Representative images of cleaved caspase-3 (green) in OC cells. (a) NT/γδ T cells, (b) 10 μM BMS202 alone, (c) ND alone, (d) ND/2.5 μM BMS202, (e) ND/5 μM BMS202, (f) ND/10 μM BMS202. Nuclei in blue (Hoechst). Scale bar = 200 μm. (g) Quantification of cleaved caspase-3 intensity for OCAS7 and (h) Quantification for OCAS8. Statistical significance was determined using one-way ANOVA followed by Dunnett’s post hoc test. The data were represented as mean ± SEM (n=3). “*” for p < 0.05, “**” for p < 0.01; and “***” for p < 0.001. vs. NT control.

**Figure 11.**
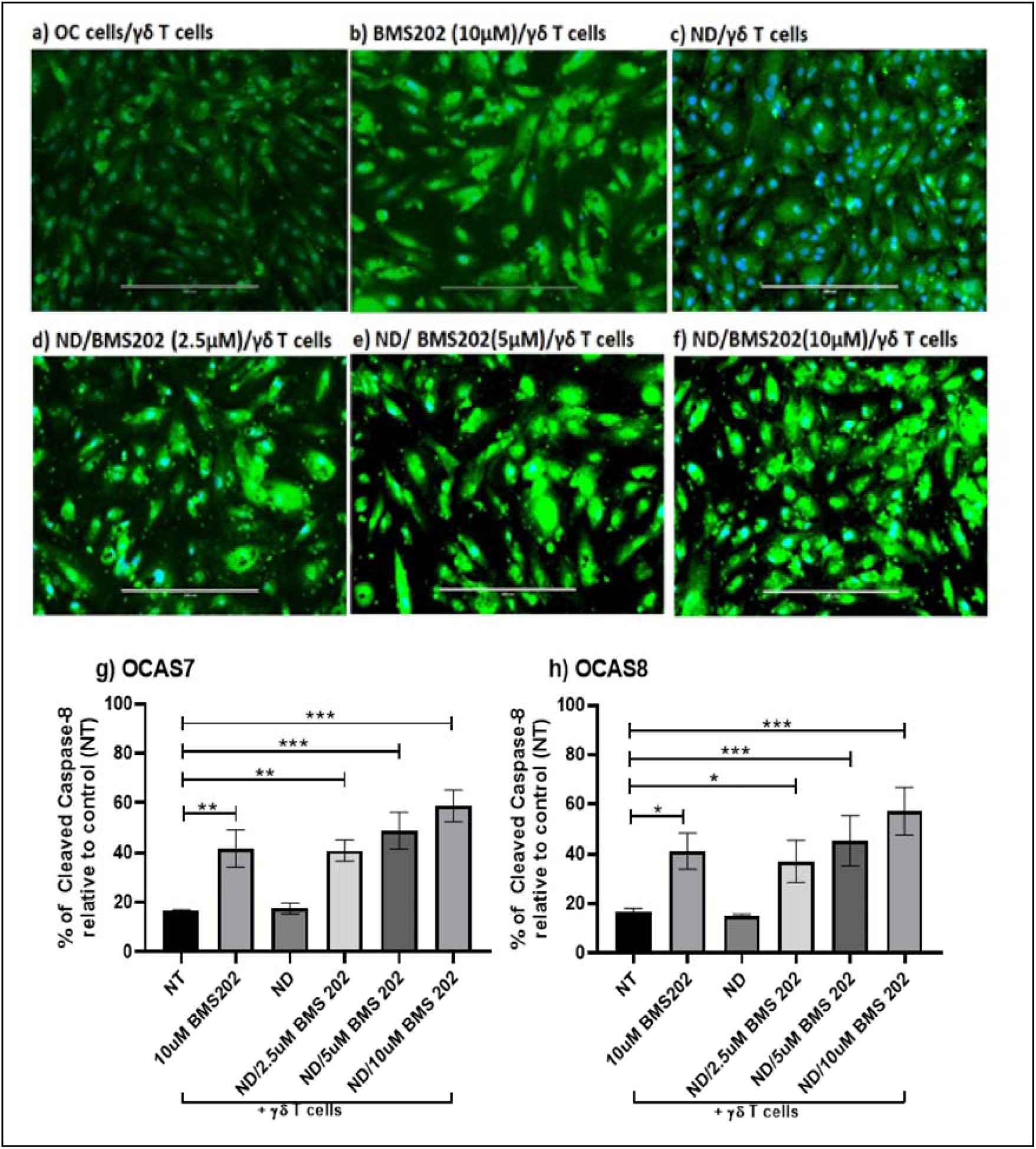
Cleaved caspase-8 expression in OC cells after ND/BMS202 and γδ T cell coculture. (a–f) Representative images of cleaved caspase-8 (green) in OC cells. (a) NT/γδ T cells, (b) 10 μM BMS202 alone, (c) ND alone, (d) ND/2.5 μM BMS202, (e) ND/5 μM BMS202, (f) ND/10 μM BMS202. Nuclei in blue (Hoechst). Scale bar = 200 μm. (g) Quantification of cleaved caspase-8 intensity for OCAS7 and (h) Quantification for OCAS8. Statistical significance was determined using one-way ANOVA followed by Dunnett’s post hoc test. The data were represented as mean ± SEM (n=3). “*” for p < 0.05, “**” for p < 0.01; and “***” for p < 0.001. vs. NT control.

Nanocomplex treatment also amplified DNA damage following γδ T-cell engagement. ND/BMS202 at 5**□**µM and 10**□**µM significantly increased γ-H2AX expression in both cell lines (p<0.001), with the 10**□**µM nanocomplex outperforming free BMS202 (p<0.01 for OCAS7; p<0.05 for OCAS8). Naked NDs did not induce γ-H2AX. Consistent with these mechanistic markers, ND/BMS202 significantly enhanced γδ T cell-mediated killing (Fig.**□**12g–h and Supplementary Tables 11 & 12). Cytell imaging revealed a marked reduction in viable cells after treatment with ND/5**□**µM or ND/10**□**µM BMS202, or with 10**□**µM free BMS202 (all p<0.001). The 2.5**□**µM nanocomplex reduced viability in OCAS7 (p<0.001) but had a weaker effect in OCAS8 (p<0.01). Naked NDs did not alter viability (Fig.**□**13 a-b and Supplementary Table 11 & 12). Together, these data demonstrate that ND/BMS202 nanocomplexes potentiate γδ T-cell cytotoxicity by activating initiator and executioner caspases, inducing substantial DNA damage, and reducing tumour cell survival in a dose-dependent manner. The ND delivery system enhances the functional impact of BMS202, producing stronger apoptotic and genotoxic responses than the free inhibitor.

**Figure 12.**
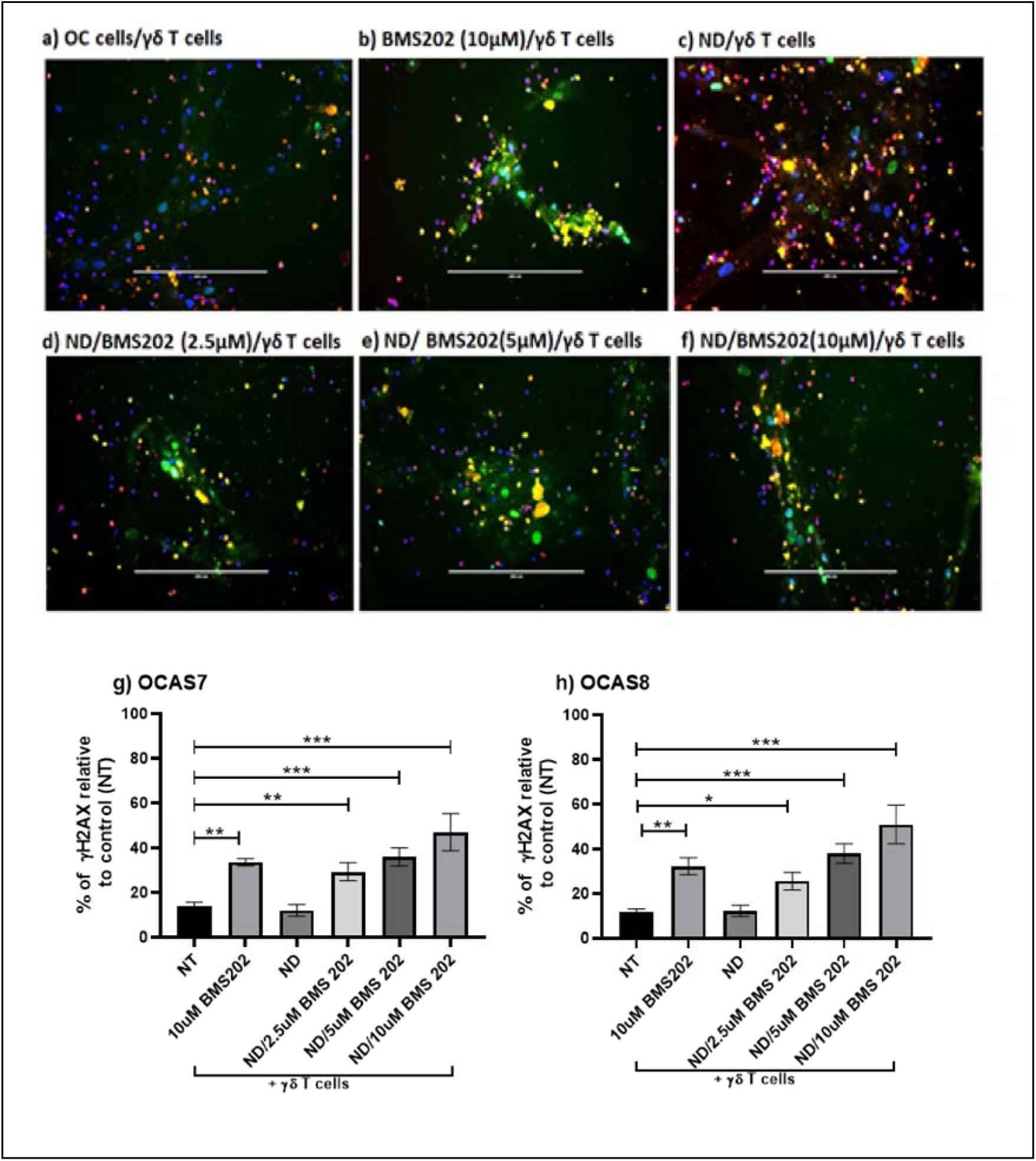
Detection of γ□H2AX nuclear expression in OC response post-exposure to ND/BMS202/ γδ T cells. (a-f) representative images of γ**□**H2AX nuclear expression in OCAS7 cells. OCAS7 and OCAS8 cells were exposed to platelets (at a ratio of 2000 platelets to 1 cancer cell) for 48 h. a) OC cells were either co-cultured with γδ T cells (NT/γδ T cells) or (b), treated with10 μM BMS202 alone (c), ND (d), ND/2.5 μM BMS202 (e), ND/5 μM BMS202 and (f) ND/10 μM BMS202 for 6 h, then cells were co-cultured with γδ T cells (10 γδ T cells to 1 cancer cell) for an additional 24 h. OC cells were probed using anti-γ**□**H2AX antibody (green) and PE-conjugated anti-CD45 antibody (red), and counterstained using Hoechst 33342). Then the expression of DNA damage marker (γ**□**H2AX) was examined using the Cytell ™ imaging system, the number of γ**□**H2AX-positive nuclei was counted (g,h) using BioApp software. Statistical significance was determined using one-way ANOVA followed by Dunnett’s post hoc test. The data were represented as mean ± SEM (n=3). “*” for p < 0.05, “**” for p < 0.01; and “***” for p < 0.001. vs. NT control.

**Figure 13.**
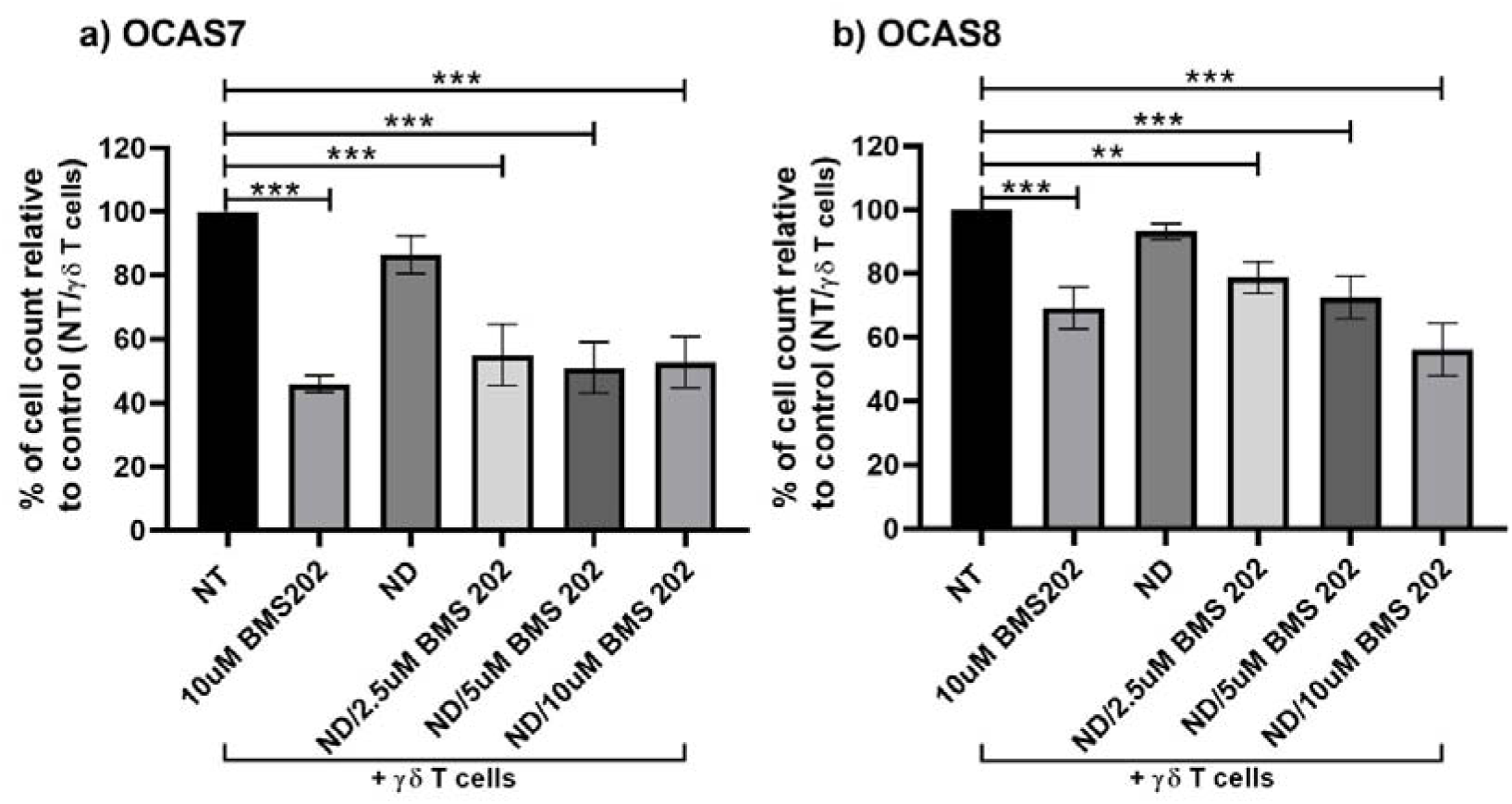
OC cell viability after ND/BMS202 and γδ T cell co-culture. Quantification of viable OCAS7 cells (a), and OCAS8 cells (b). Statistical significance was determined using one-way ANOVA followed by Dunnett’s post hoc test. The data were represented as mean ± SEM (n=3). “*” for p < 0.05, “**” for p < 0.01; and “***” for p < 0.001. vs. NT control.

## Discussion

Ovarian cancer remains the deadliest gynaecological malignancy, largely because of late diagnosis, high metastatic potential, rapid dissemination within the peritoneal cavity, and frequent acquisition of resistance to therapy (1–5,41–44). γδ T cells possess strong cytotoxic potential against metastatic and stem-like OC cells, but their activity can be limited by immunosuppressive factors within the tumour microenvironment, including PD-L1 expression and physical barriers such as platelet cloaking (45–50). This study evaluated a combination strategy integrating ex vivo expanded γδ T cells with an ND-delivered PD-L1 inhibitor (BMS202) and demonstrated enhanced γδ T-cell cytotoxicity against platelet-cloaked OC cells.

We first established clinically relevant models using ascites-derived OC cell cultures (OCAS7 and OCAS8), showing the expression of PAX8, WT1, and p16, to support the conclusions on ovarian origin (36–38,51,52). To model metastatic adaption in the circulation, the OC cells were cloaked with platelets, which induced morphological changes consistent with an epithelial to mesenchymal transition. Increased N-cadherin, β-catenin and thrombin expression, together with reduced E-cadherin expression, supported this phenotypic shift (10,53–60). Platelets are known to directly facilitate immune escape, increase CTC survival and promoting metastatic cancer invasion (54,58,61–64), indicating that this OC cell model effectively recapitulates key features of disseminated advanced disease.

The expanded γδ T-cell population contained Vδ1**□** (24%), Vδ2**□** (14.3%), and Vδ3**□** (9.5%) subsets, each of which may contribute distinct modes of tumour recognition (20,21,25,45–48). Vδ1**□** T cells recognise stress-induced ligands such as MICA/B and ULBPs; Vδ2**□** T cells recognise phosphoantigens associated with malignant transformation; and Vδ3**□** T cells recognise lipids presented by CD1d. γδ T cells have been identified in OC ascites, where their abundance has been associated with improved patient outcomes (47,48). The MHC-independent recognition of cancer cells by γδ T cells, together with rapid effector function, direct cytotoxicity, and ability to infiltrate hypoxic tumour regions make these T cells particularly attractive candidates for solid tumour immunotherapy (20,46,65). However the potential for metastatic tumour cells to negate the cytotoxic action of γδ T cells by upregulating the surface expression of PD-1 to bind PDL-1 and induce immune suppression is a significant limitation for therapeutic efficacy.

The ND/BMS202 nanocomplex may facilitate targeted PD**□**1/PD**□**L1 inhibition to significantly enhance γδ T-cell activation and OC cell toxicity. This ND-based delivery system offers significant advantages including improved targeting, reduced off-target toxicity, and prolonged drug persistence at the tumour/immune T cell interface (29–31). ND/BMS202 increased γδ T-cell clustering and attachment to OC cells facilitating interactions that are essential for cytotoxicity, immune synapse formation, and the disruption of PD-1/PD-L1 signalling (22,25–27). This enhanced contact was accompanied by increased degranulation (CD107a) and GrB release (22,26,65), which are direct measures of cytotoxic T-cell action. The nanocomplex may therefore exert a dual activity: BMS202 negating PD-1/PD-L1 interactions, while the ND scaffold may facilitate immune synapse formation. CD107a and GrB are well-established correlates of cytotoxic function (39,66–68), and the nanocomplex outperformed free BMS202 in a dose-dependent manner for OC cell cytotoxicity.

Mechanistic analyses demonstrated activation of both extrinsic and granzyme-mediated apoptotic pathways, supporting the conclusions on increased cytotoxicity. Cleaved caspase-8 and cleaved caspase-3 were significantly upregulated following treatment, consistent with FasL/TRAIL signalling and GrB-mediated apoptosis (39,66,68). Strong γ-H2AX staining indicated extensive DNA double-strand breaks and damage to the OC cells (40,69–71). Nanocomplex-treated cells exhibited significantly greater DNA damage than those treated with free BMS202 despite the presence of platelet cloaks, confirming efficient nanoparticle-mediated delivery. Similar enhancements have been reported with other nanoparticle platforms (32,72). Although free BMS202 retained activity, the ND-conjugated formulation demonstrated superior potency, particularly at lower doses likely due to enhanced uptake and sustained release (31,32,73–76). Naked NDs did not alter GrB release, caspase activation, or viability, confirming that the observed effects were attributable to PD-L1 inhibition rather than the carrier (73–76).

These findings align with emerging directions in OC immunotherapy, reinforcing the potential of γδ T cells as versatile effector cells capable of functioning within the peritoneal environment (47,50). The findings also highlight the importance of targeting additional immunosuppressive components of the tumour microenvironment, such as tumour-associated macrophages, and other immune cells that drive EMT and therapeutic resistance (9,49). Nanomedicine-based strategies are well positioned to address these challenges, and prior work has shown that ND-delivered BMS202 can augment immune activity in melanoma models (32,77,78). The present findings extend this concept to OC, demonstrating that ND-mediated delivery enhances PD-L1 inhibition and amplifies γδ T-cell effector functions, including degranulation and GrB-dependent cytotoxicity.

Several limitations should be acknowledged. The system used here is two-dimensional and does not capture the complex cellular interactions and physical constraints of the *in vivo* tumour microenvironment. The OC microenvironment includes fibroblasts, macrophages, regulatory T cells, and soluble mediators that were not incorporated into the current model (49,50). Patient derived organoids (PDOs) represent a more physiologically relevant platform for the next stage of validation, preserving tumour architecture and genetic diversity, and have already been used to assess immune-mediated responses in OC (45,79–81). Additionally, the platelet-cloaking model, while informative, remains an *in vitro* approximation of peritoneal dynamics (10,53). Future work should therefore evaluate the ND/BMS202 platform in 3D/PDO systems and immunocompetent animal models. Interactions between platelet cloaking, nanocomplex delivery, and tumour-associated macrophages warrant further investigation, as they may reveal new combination strategies. Engineering γδ CAR T cells represents another promising direction, potentially enabling off-the-shelf cell therapy (20,26,28). Pharmacokinetic, biodistribution and target-engagement studies are also required to confirm tumour accumulation of NDs, define BMS202 delivery kinetics and assess safety (29,31).

In summary, nanoparticle-mediated delivery of a PD-L1 inhibitor enhanced γδ T-cell efficacy against metastatic platelet-cloaked OC cells. The ND/BMS202 nanocomplex improved T-cell–tumour contact, degranulation, GrB production, apoptotic signalling, DNA damage and tumour-cell killing across two patient-derived OC cell lines.. These findings provide proof-of-principle for further therapeutic validation of ND/BMS202 as part of a γδ T-cell-based combination strategy. Integrating nanomedicine, γδ T-cell biology and advanced tumour models offers a promising route toward more effective immunotherapeutic approaches for this highly lethal disease.

## Supplemental Information

**Table S1.** Comparison of GrB expression levels between different treatment groups of OCAS7 cells post treatment either BMS202 alone or with three concentrations (2,5, 5 and 10uM) of our lab developed NCs and then co cultured with γδ T cells. One-way ANOVA with Tukey’s post hoc test was used for all pairwise comparisons between treatment groups (BMS202 + NCs at 2.5, 5, or 10 µM vs. free BMS202 alone vs. NCs alone vs. NT control). Data are presented as mean differences with 95% confidence intervals (CI).

| **Tukey's multiple comparisons test** | **Mean Diff.** | **95.00% CI of diff.** | **Significant?** | **Summary** |
| --- | --- | --- | --- | --- |
| NT/$\gamma\delta$ T cells vs. 10uM BMS202/$\gamma\delta$ T cells | -0.2608 | -0.4594 to -0.06230 | Yes | ** |
| NT/$\gamma\delta$ T cells vs. ND/$\gamma\delta$ T cells | -0.1315 | -0.3300 to 0.06704 | No | ns |
| NT/$\gamma\delta$ T cells vs. ND/2.5uM BMS202/$\gamma\delta$ T cells | -0.3106 | -0.5091 to -0.1120 | Yes | ** |
| NT/$\gamma\delta$ T cells vs. ND/5uM BMS202/$\gamma\delta$ T cells | -0.3869 | -0.5855 to -0.1884 | Yes | *** |
| NT/$\gamma\delta$ T cells vs. ND/10uM BMS202/$\gamma\delta$ T cells | -0.4573 | -0.6558 to -0.2587 | Yes | *** |
| 10uM BMS202/$\gamma\delta$ T cells vs. ND/$\gamma\delta$ T cells | 0.1293 | -0.06919 to 0.3279 | No | ns |
| 10uM BMS202/$\gamma\delta$ T cells vs. ND/2.5uM BMS202/$\gamma\delta$ T cells | -0.04973 | -0.2483 to 0.1488 | No | ns |
| 10uM BMS202/$\gamma\delta$ T cells vs. ND/5uM BMS202/$\gamma\delta$ T cells | -0.1261 | -0.3246 to 0.07245 | No | ns |
| 10uM BMS202/$\gamma\delta$ T cells vs. ND/10uM BMS 202/$\gamma\delta$ T cells | -0.1964 | -0.3950 to 0.002099 | No | ns |
| ND/$\gamma\delta$ T cells vs. ND/2.5uM BMS202/$\gamma\delta$ T cells | -0.1791 | -0.3776 to 0.01946 | No | ns |
| ND/$\gamma\delta$ T cells vs. ND/5uM BMS 202/$\gamma\delta$ T cells | -0.2554 | -0.4540 to -0.05689 | Yes | ** |
| ND/$\gamma\delta$ T cells vs. ND/10uM BMS202/$\gamma\delta$ T cells | -0.3258 | -0.5243 to -0.1272 | Yes | ** |
| ND/2.5uM BMS202/$\gamma\delta$ T cells vs. ND/5uM BMS202/$\gamma\delta$ T cells | -0.07635 | -0.2749 to 0.1222 | No | ns |
| ND/2.5uM BMS202/$\gamma\delta$ T cells vs. ND/10uM BMS202/$\gamma\delta$ T cells | -0.1467 | -0.3452 to 0.05183 | No | ns |

**Table S2.** Comparison of GrB expression levels between different treatment groups of OCAS8 cells post treatment either BMS202 alone or with three concentrations (2,5, 5 and 10uM) of our lab developed NCs and then co cultured with γδ T cells. One-way ANOVA with Tukey’s post hoc test was used for all pairwise comparisons between treatment groups (BMS202 + NCs at 2.5, 5, or 10 µM vs. free BMS202 alone vs. NCs alone vs. NT control). Data are presented as mean differences with 95% confidence intervals (CI).

| **Tukey's multiple comparisons test** | **Mean Diff.** | **95.00% CI of diff.** | **Significant?** | **Summary** |
| --- | --- | --- | --- | --- |
| NT/$\gamma\delta$ T cells vs. 10uM BMS202/$\gamma\delta$ T cells | -0.2942 | -0.5367 to -0.05168 | Yes | * |
| NT/$\gamma\delta$ T cells vs. ND/$\gamma\delta$ T cells | -0.1648 | -0.4073 to 0.07767 | No | ns |
| NT/$\gamma\delta$ T cells vs. ND/2.5uM BMS202/$\gamma\delta$ T cells | -0.2772 | -0.5197 to -0.03474 | Yes | * |
| NT/$\gamma\delta$ T cells vs. ND/5uM BMS202/$\gamma\delta$ T cells | -0.4869 | -0.7294 to -0.2444 | Yes | *** |
| NT/$\gamma\delta$ T cells vs. ND/10uM BMS202/$\gamma\delta$ T cells | -0.5906 | -0.8331 to -0.3481 | Yes | *** |
| 10uM BMS202/$\gamma\delta$ T cells vs. ND/$\gamma\delta$ T cells | 0.1293 | -0.1132 to 0.3718 | No | ns |
| 10uM BMS202/$\gamma\delta$ T cells vs. ND/2.5uM BMS202/$\gamma\delta$ T cells | 0.01693 | -0.2256 to 0.2594 | No | ns |
| 10uM BMS202/$\gamma\delta$ T cells vs. ND/5uM BMS202/$\gamma\delta$ T cells | -0.1928 | -0.4352 to 0.04974 | No | ns |
| 10uM BMS202/$\gamma\delta$ T cells vs. ND/10uM BMS 202/$\gamma\delta$ T cells | -0.2964 | -0.5389 to -0.05394 | Yes | * |
| ND/$\gamma\delta$ T cells vs. ND/2.5uM BMS202/$\gamma\delta$ T cells | -0.1124 | -0.3549 to 0.1301 | No | ns |
| ND/$\gamma\delta$ T cells vs. ND/5uM BMS 202/$\gamma\delta$ T cells | -0.3221 | -0.5646 to -0.07960 | Yes | ** |
| ND/$\gamma\delta$ T cells vs. ND/10uM BMS202/$\gamma\delta$ T cells | -0.4258 | -0.6683 to -0.1833 | Yes | *** |
| ND/2.5uM BMS202/$\gamma\delta$ T cells vs. ND/5uM BMS202/$\gamma\delta$ T cells | -0.2097 | -0.4522 to 0.03281 | No | ns |
| ND/2.5uM BMS202/$\gamma\delta$ T cells vs. ND/10uM BMS202/$\gamma\delta$ T cells | -0.3134 | -0.5559 to -0.07088 | Yes | ** |

**Table S3.**
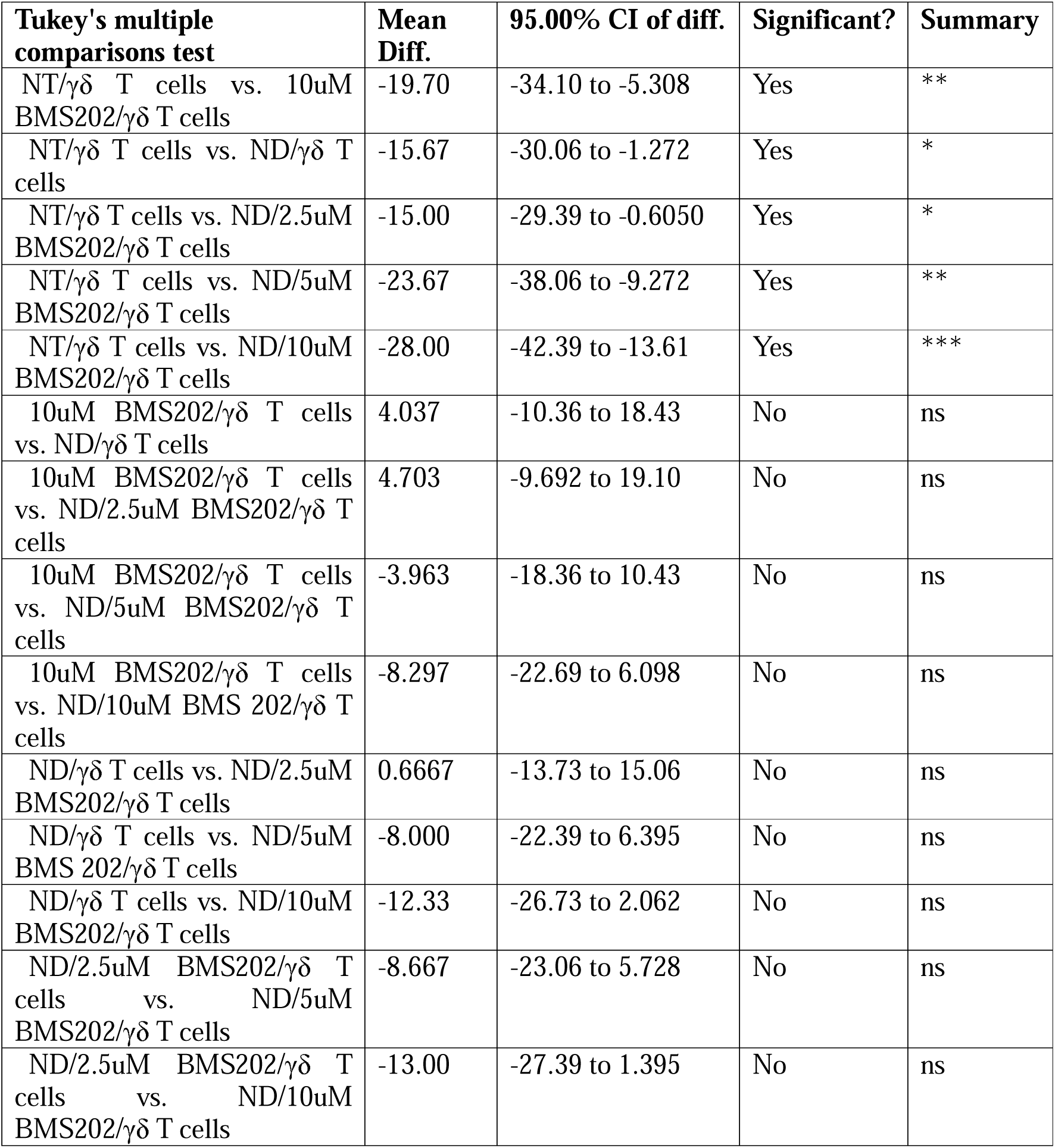
Comparison of CD107a degranulation levels between different treatment groups of OCAS7 cells post treatment either BMS202 alone or with three concentrations (2,5, 5 and 10uM) of our lab developed NCs and then co cultured with γδ T cells. One-way ANOVA with Tukey’s post hoc test was used for all pairwise comparisons between treatment groups (BMS202 + NCs at 2.5, 5, or 10 µM vs. free BMS202 alone vs. NCs alone vs. NT control). Data are presented as mean differences with 95% confidence intervals (CI).

**Table S4.**
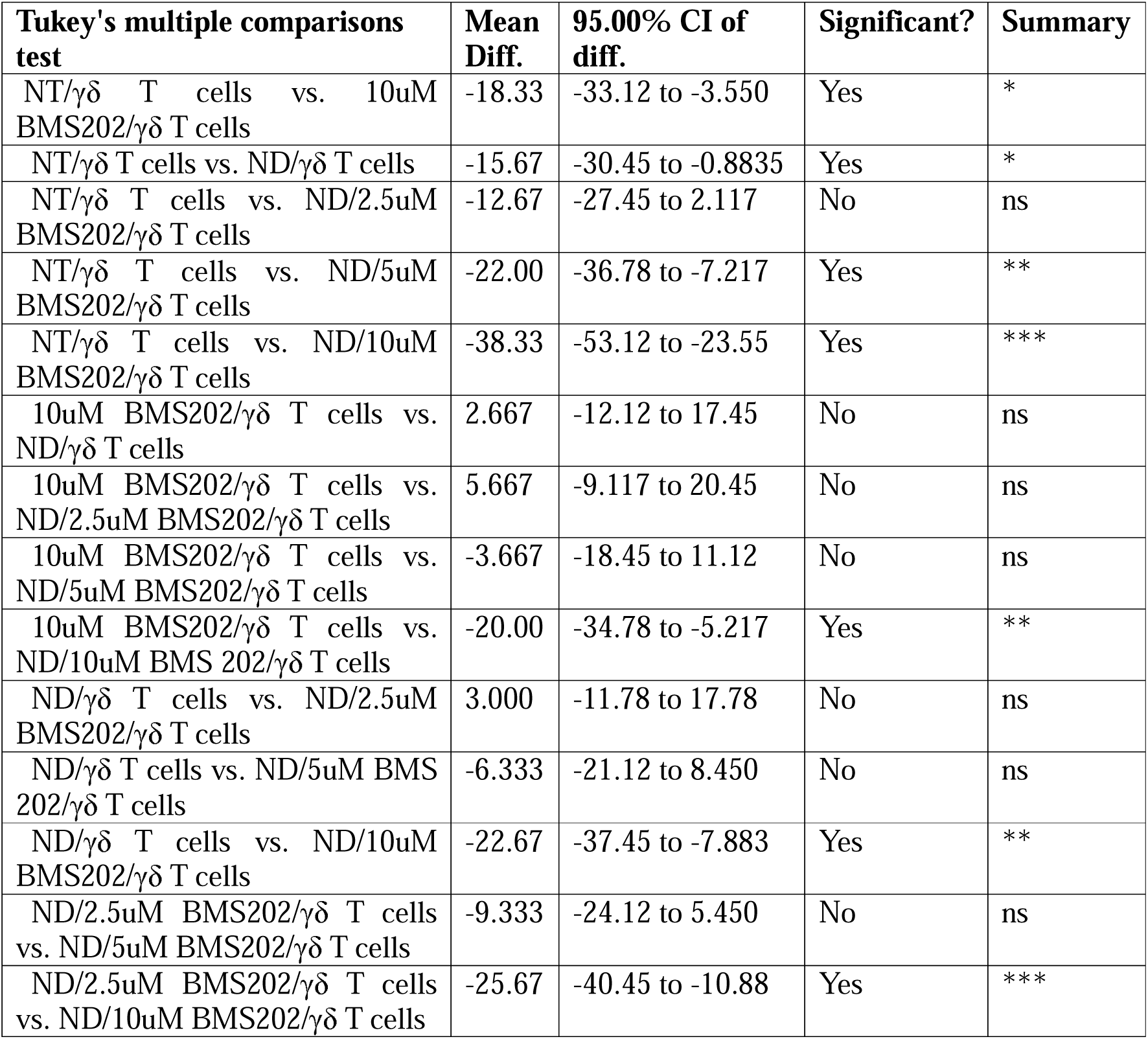
Comparison of CD107a degranulation levels between different treatment groups of OCAS8 cells post treatment either BMS202 alone or with three concentrations (2,5, 5 and 10uM) of our lab developed NCs and then co cultured with γδ T cells. One-way ANOVA with Tukey’s post hoc test was used for all pairwise comparisons between treatment groups (BMS202 + NCs at 2.5, 5, or 10 µM vs. free BMS202 alone vs. NCs alone vs. NT control). Data are presented as mean differences with 95% confidence intervals (CI).

**Table S5.**
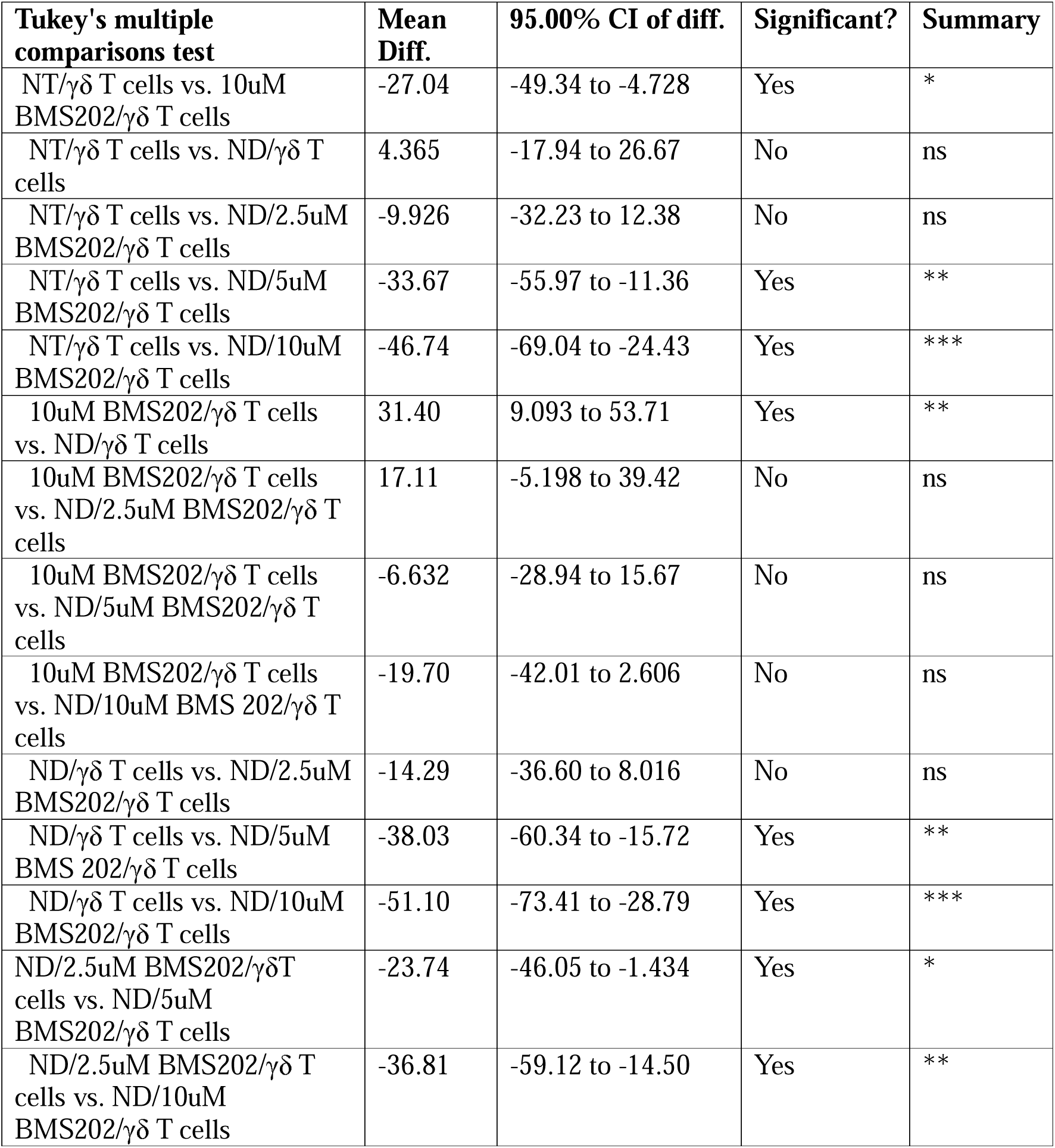
Comparison of cleaved caspase 3 expression levels between different treatment groups of OCAS7 cells post treatment either BMS202 alone or with three concentrations (2,5, 5 and 10uM) of our lab developed NCs and then co cultured with γδ T cells. One-way ANOVA with Tukey’s post hoc test was used for all pairwise comparisons between treatment groups (BMS202 + NCs at 2.5, 5, or 10 µM vs. free BMS202 alone vs. NCs alone vs. NT control). Data are presented as mean differences with 95% confidence intervals (CI).

**Table S6.**
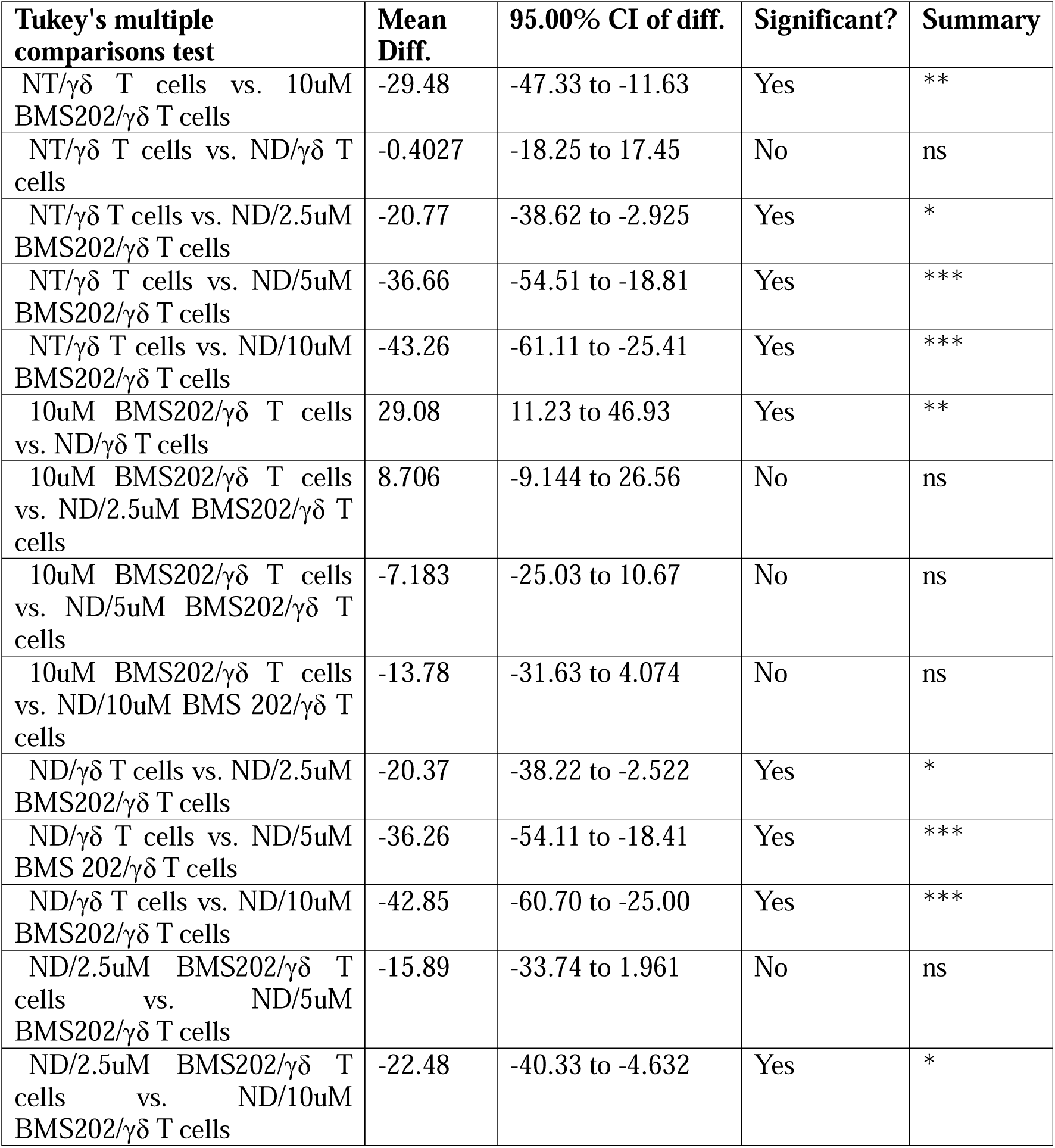
Comparison of cleaved caspase 3 expression levels between different treatment groups of OCAS8 cells post treatment either BMS202 alone or with three concentrations (2,5, 5 and 10uM) of our lab developed NCs and then co cultured with γδ T cells. One-way ANOVA with Tukey’s post hoc test was used for all pairwise comparisons between treatment groups (BMS202 + NCs at 2.5, 5, or 10 µM vs. free BMS202 alone vs. NCs alone vs. NT control). Data are presented as mean differences with 95% confidence intervals (CI).

**Table S7.**
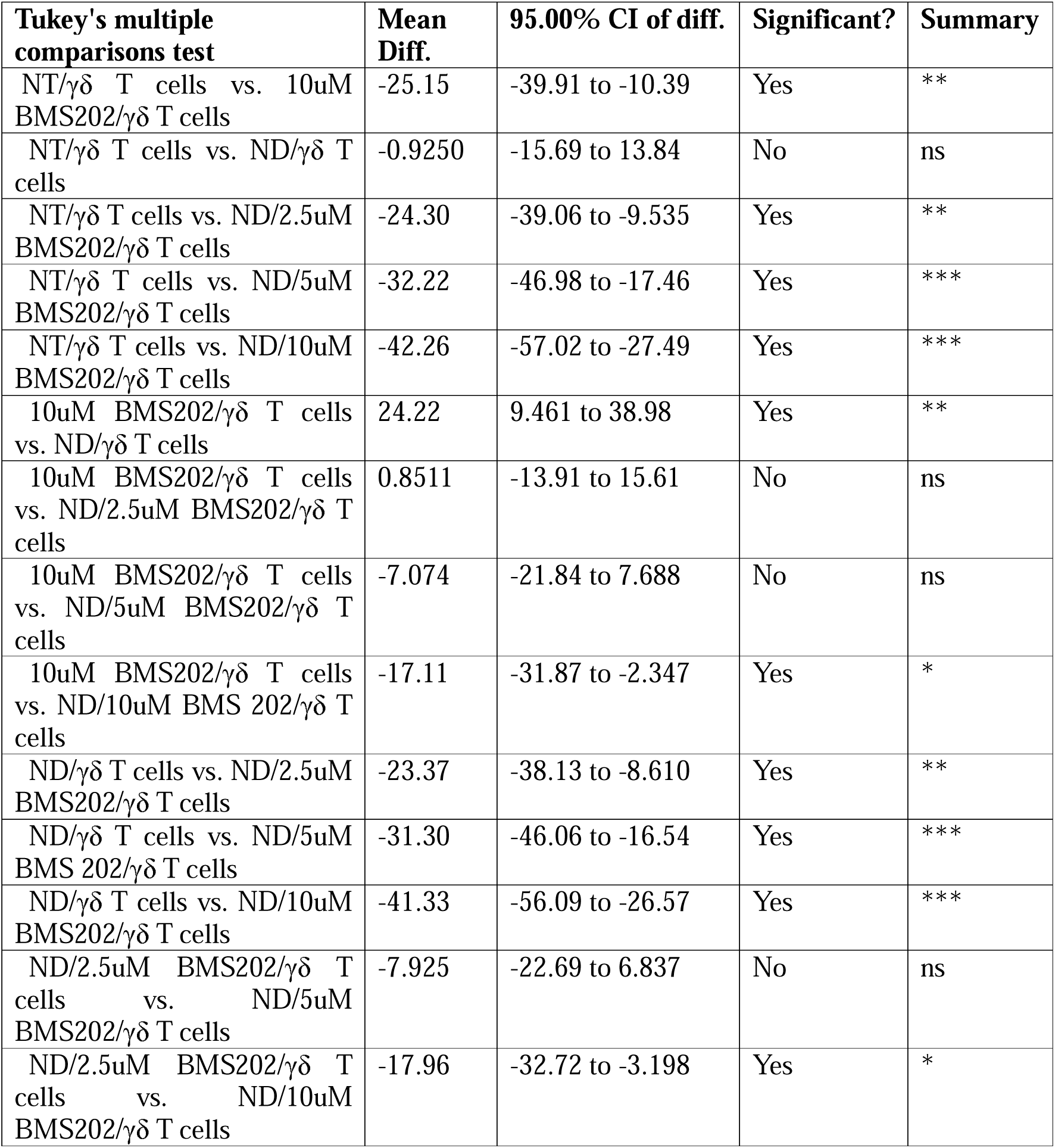
Comparison of cleaved caspase 8 expression levels between different treatment groups of OCAS7 cells post treatment either BMS202 alone or with three concentrations (2,5, 5 and 10uM) of our lab developed NCs and then co cultured with γδ T cells. . One-way ANOVA with Tukey’s post hoc test was used for all pairwise comparisons between treatment groups (BMS202 + NCs at 2.5, 5, or 10 µM vs. free BMS202 alone vs. NCs alone vs. NT control). Data are presented as mean differences with 95% confidence intervals (CI).

**Table S8.**
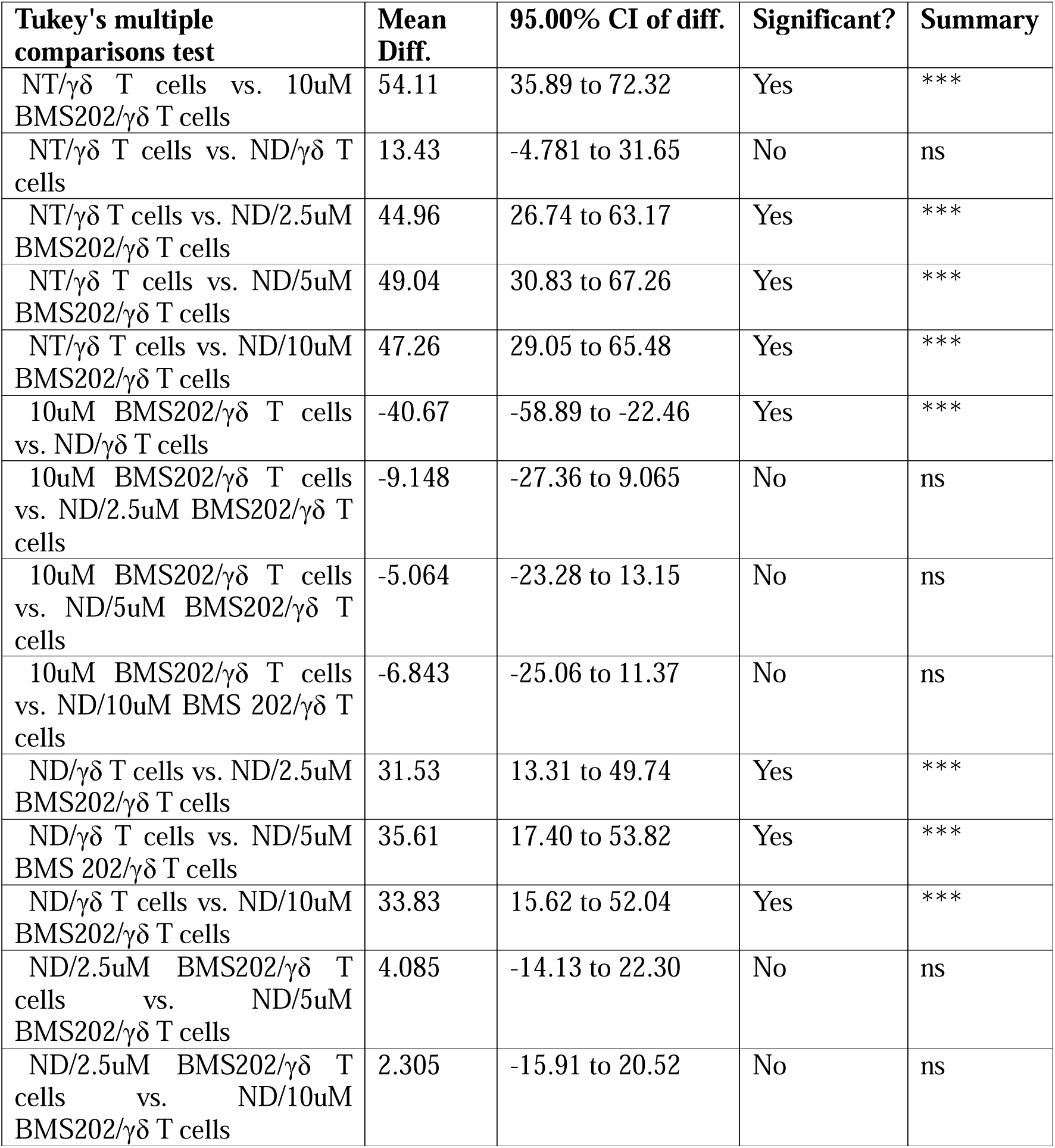
Comparison of cleaved caspase 8 expression levels between different treatment groups of OCAS8 cells post treatment either BMS202 alone or with three concentrations (2,5, 5 and 10uM) of our lab developed NCs and then co cultured with γδ T cells. One-way ANOVA with Tukey’s post hoc test was used for all pairwise comparisons between treatment groups (BMS202 + NCs at 2.5, 5, or 10 µM vs. free BMS202 alone vs. NCs alone vs. NT control). Data are presented as mean differences with 95% confidence intervals (CI).

| **Tukey's multiple comparisons test** | **Mean Diff.** | **95.00% CI of diff.** | **Significant?** | **Summary** |
| --- | --- | --- | --- | --- |
| NT/$\gamma\delta$ T cells vs. 10uM BMS202/$\gamma\delta$ T cells | 54.11 | 35.89 to 72.32 | Yes | *** |
| NT/$\gamma\delta$ T cells vs. ND/$\gamma\delta$ T cells | 13.43 | -4.781 to 31.65 | No | ns |
| NT/$\gamma\delta$ T cells vs. ND/2.5uM BMS202/$\gamma\delta$ T cells | 44.96 | 26.74 to 63.17 | Yes | *** |
| NT/$\gamma\delta$ T cells vs. ND/5uM BMS202/$\gamma\delta$ T cells | 49.04 | 30.83 to 67.26 | Yes | *** |
| NT/$\gamma\delta$ T cells vs. ND/10uM BMS202/$\gamma\delta$ T cells | 47.26 | 29.05 to 65.48 | Yes | *** |
| 10uM BMS202/$\gamma\delta$ T cells vs. ND/$\gamma\delta$ T cells | -40.67 | -58.89 to -22.46 | Yes | *** |
| 10uM BMS202/$\gamma\delta$ T cells vs. ND/2.5uM BMS202/$\gamma\delta$ T cells | -9.148 | -27.36 to 9.065 | No | ns |
| 10uM BMS202/$\gamma\delta$ T cells vs. ND/5uM BMS202/$\gamma\delta$ T cells | -5.064 | -23.28 to 13.15 | No | ns |
| 10uM BMS202/$\gamma\delta$ T cells vs. ND/10uM BMS 202/$\gamma\delta$ T cells | -6.843 | -25.06 to 11.37 | No | ns |
| ND/$\gamma\delta$ T cells vs. ND/2.5uM BMS202/$\gamma\delta$ T cells | 31.53 | 13.31 to 49.74 | Yes | *** |
| ND/$\gamma\delta$ T cells vs. ND/5uM BMS 202/$\gamma\delta$ T cells | 35.61 | 17.40 to 53.82 | Yes | *** |
| ND/$\gamma\delta$ T cells vs. ND/10uM BMS202/$\gamma\delta$ T cells | 33.83 | 15.62 to 52.04 | Yes | *** |
| ND/2.5uM BMS202/$\gamma\delta$ T cells vs. ND/5uM BMS202/$\gamma\delta$ T cells | 4.085 | -14.13 to 22.30 | No | ns |
| ND/2.5uM BMS202/$\gamma\delta$ T cells vs. ND/10uM BMS202/$\gamma\delta$ T cells | 2.305 | -15.91 to 20.52 | No | ns |

**Table S9.** Comparison of cell. γ **H2AX expression between different treatment groups of OCAS7 cells post treatment either BMS202 alone or with three concentrations (2,5, 5 and 10uM) of our lab developed NCs and then co cultured with γδ T cells.** One-way ANOVA with Tukey’s post hoc test was used for all pairwise comparisons between treatment groups (BMS202 + NCs at 2.5, 5, or 10 µM vs. free BMS202 alone vs. NCs alone vs. NT control). Data are presented as mean differences with 95% confidence intervals (CI).

| **Tukey's multiple comparisons test** | **Mean Diff.** | **95.00% CI of diff.** | **Significant?** | **Summary** |
| --- | --- | --- | --- | --- |
| NT/$\gamma\delta$ T cells vs. 10uM BMS202/$\gamma\delta$ T cells | -20.00 | -31.90 to -8.098 | Yes | ** |
| NT/$\gamma\delta$ T cells vs. ND/$\gamma\delta$ T cells | 1.667 | -10.24 to 13.57 | No | ns |
| NT/$\gamma\delta$ T cells vs. ND/2.5uM BMS202/$\gamma\delta$ T cells | -15.67 | -27.57 to -3.765 | Yes | ** |
| NT/$\gamma\delta$ T cells vs. ND/5uM BMS202/$\gamma\delta$ T cells | -22.33 | -34.24 to -10.43 | Yes | *** |
| NT/$\gamma\delta$ T cells vs. ND/10uM BMS202/$\gamma\delta$ T cells | -33.33 | -45.24 to -21.43 | Yes | *** |
| 10uM BMS202/$\gamma\delta$ T cells vs. ND/$\gamma\delta$ T cells | 21.67 | 9.765 to 33.57 | Yes | *** |
| 10uM BMS202/$\gamma\delta$ T cells vs. ND/2.5uM BMS202/$\gamma\delta$ T cells | 4.333 | -7.569 to 16.24 | No | ns |
| 10uM BMS202/$\gamma\delta$ T cells vs. ND/5uM BMS202/$\gamma\delta$ T cells | -2.333 | -14.24 to 9.569 | No | ns |
| 10uM BMS202/$\gamma\delta$ T cells vs. ND/10uM BMS 202/$\gamma\delta$ T cells | -13.33 | -25.24 to -1.431 | Yes | * |
| ND/$\gamma\delta$ T cells vs. ND/2.5uM BMS202/$\gamma\delta$ T cells | -17.33 | -29.24 to -5.431 | Yes | ** |
| ND/$\gamma\delta$ T cells vs. ND/5uM BMS 202/$\gamma\delta$ T cells | -24.00 | -35.90 to -12.10 | Yes | *** |
| ND/$\gamma\delta$ T cells vs. ND/10uM BMS202/$\gamma\delta$ T cells | -35.00 | -46.90 to -23.10 | Yes | *** |
| ND/2.5uM BMS202/$\gamma\delta$ T cells vs. ND/5uM BMS202/$\gamma\delta$ T cells | -6.667 | -18.57 to 5.235 | No | ns |
| ND/2.5uM BMS202/$\gamma\delta$ T cells vs. ND/10uM BMS202/$\gamma\delta$ T cells | -17.67 | -29.57 to -5.765 | Yes | ** |

**Table S10.** Comparison of cell. γ **H2AX expression between different treatment groups of OCAS8 cells post treatment either BMS202 alone or with three concentrations (2,5, 5 and 10uM) of our lab developed NCs and then co cultured with γδ T cells.** One-way ANOVA with Tukey’s post hoc test was used for all pairwise comparisons between treatment groups (BMS202 + NCs at 2.5, 5, or 10 µM vs. free BMS202 alone vs. NCs alone vs. NT control). Data are presented as mean differences with 95% confidence intervals (CI).

| **Tukey's multiple comparisons test** | **Mean Diff.** | **95.00% CI of diff.** | **Significant?** | **Summary** |
| --- | --- | --- | --- | --- |
| NT/$\gamma\delta$ T cells vs. 10uM BMS202/$\gamma\delta$ T cells | -20.67 | -33.50 to -7.836 | Yes | ** |
| NT/$\gamma\delta$ T cells vs. ND/$\gamma\delta$ T cells | -0.6667 | -13.50 to 12.16 | No | ns |
| NT/$\gamma\delta$ T cells vs. ND/2.5uM BMS202/$\gamma\delta$ T cells | -14.00 | -26.83 to -1.169 | Yes | * |
| NT/$\gamma\delta$ T cells vs. ND/5uM BMS202/$\gamma\delta$ T cells | -26.33 | -39.16 to -13.50 | Yes | *** |
| NT/$\gamma\delta$ T cells vs. ND/10uM BMS202/$\gamma\delta$ T cells | -39.33 | -52.16 to -26.50 | Yes | *** |
| 10uM BMS202/$\gamma\delta$ T cells vs. ND/$\gamma\delta$ T cells | 20.00 | 7.169 to 32.83 | Yes | ** |
| 10uM BMS202/$\gamma\delta$ T cells vs. ND/2.5uM BMS202/$\gamma\delta$ T cells | 6.667 | -6.164 to 19.50 | No | ns |
| 10uM BMS202/$\gamma\delta$ T cells vs. ND/5uM BMS202/$\gamma\delta$ T cells | -5.667 | -18.50 to 7.164 | No | ns |
| 10uM BMS202/$\gamma\delta$ T cells vs. ND/10uM BMS 202/$\gamma\delta$ T cells | -18.67 | -31.50 to -5.836 | Yes | ** |
| ND/$\gamma\delta$ T cells vs. ND/2.5uM BMS202/$\gamma\delta$ T cells | -13.33 | -26.16 to -0.5022 | Yes | * |
| ND/$\gamma\delta$ T cells vs. ND/5uM BMS 202/$\gamma\delta$ T cells | -25.67 | -38.50 to -12.84 | Yes | *** |
| ND/$\gamma\delta$ T cells vs. ND/10uM BMS202/$\gamma\delta$ T cells | -38.67 | -51.50 to -25.84 | Yes | *** |
| ND/2.5uM BMS202/$\gamma\delta$ T cells vs. ND/5uM BMS202/$\gamma\delta$ T cells | -12.33 | -25.16 to 0.4978 | No | ns |
| ND/2.5uM BMS202/$\gamma\delta$ T cells vs. ND/10uM BMS202/$\gamma\delta$ T cells | -25.33 | -38.16 to -12.50 | Yes | *** |

**Table S11.**
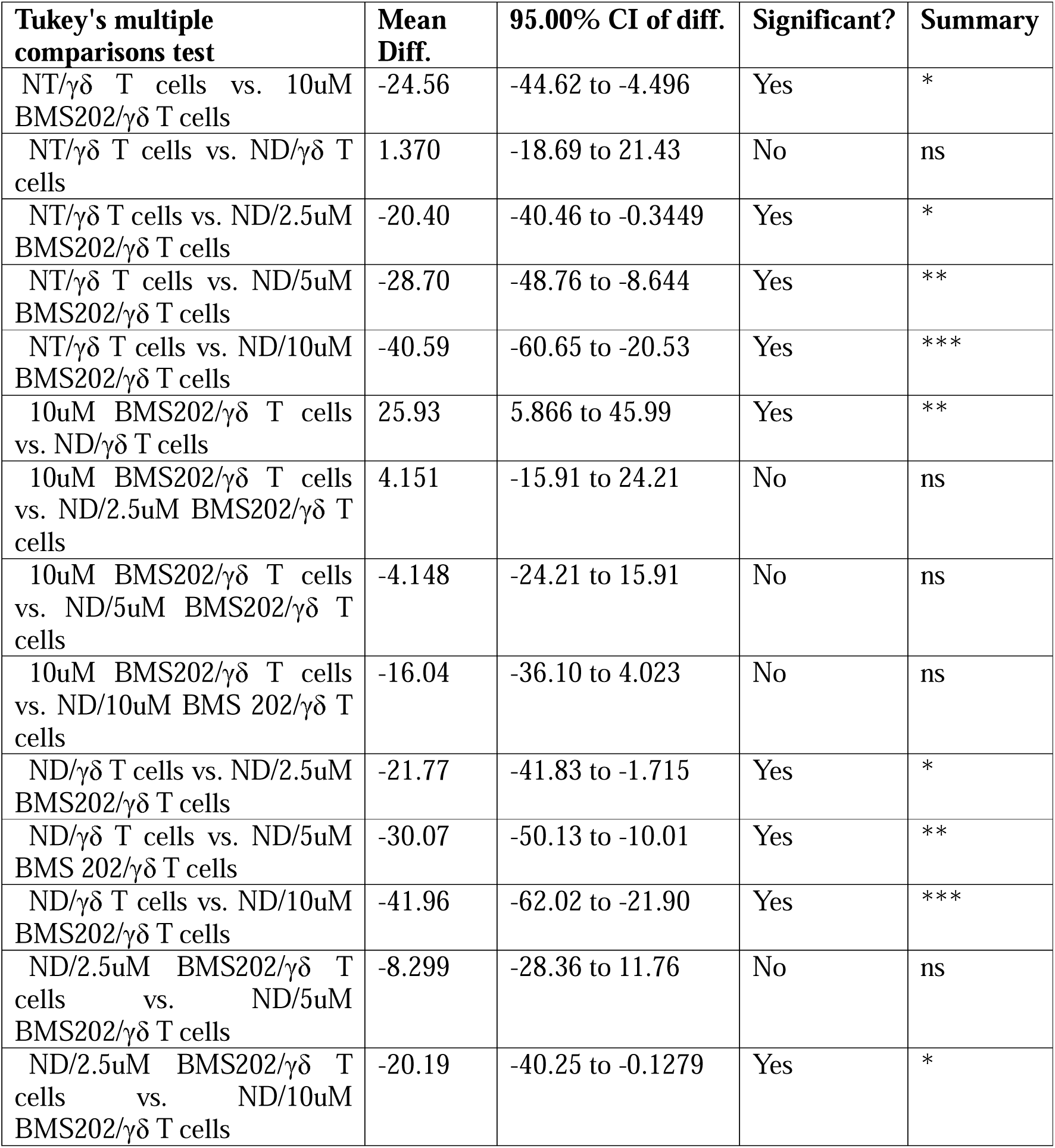
Comparison of cell viability between different treatment groups of OCAS 7 cells post treatment either BMS202 alone or with three concentrations (2,5, 5 and 10uM) of our lab developed NCs and then co cultured with γδ T cells. One-way ANOVA with Tukey’s post hoc test was used for all pairwise comparisons between treatment groups (BMS202 + NCs at 2.5, 5, or 10 µM vs. free BMS202 alone vs. NCs alone vs. NT control). Data are presented as mean differences with 95% confidence intervals (CI).

**Table S12.** Comparison of cell viability status between different treatment groups of OCAS8 Cells post treatment either BMS202 alone or with three concentrations (2,5, 5 and 10uM) of our lab developed NCs and then co cultured with γδ T cells. One-way ANOVA with Tukey’s post hoc test was used for all pairwise comparisons between treatment groups (BMS202 + NCs at 2.5, 5, or 10 µM vs. free BMS202 alone vs. NCs alone vs. NT control). Data are presented as mean differences with 95% confidence intervals (CI).

| **Tukey's multiple comparisons test** | **Mean Diff.** | **95.00% CI of diff.** | **Significant?** | **Summary** |
| --- | --- | --- | --- | --- |
| NT/$\gamma\delta$ T cells vs. 10uM BMS202/$\gamma\delta$ T cells | 30.77 | 15.56 to 45.99 | Yes | *** |
| NT/$\gamma\delta$ T cells vs. ND/$\gamma\delta$ T cells | 6.766 | -8.452 to 21.98 | No | ns |
| NT/$\gamma\delta$ T cells vs. ND/2.5uM BMS202/$\gamma\delta$ T cells | 21.29 | 6.074 to 36.51 | Yes | ** |
| NT/$\gamma\delta$ T cells vs. ND/5uM BMS202/$\gamma\delta$ T cells | 27.38 | 12.16 to 42.59 | Yes | *** |
| NT/$\gamma\delta$ T cells vs. ND/10uM BMS202/$\gamma\delta$ T cells | 43.93 | 28.71 to 59.15 | Yes | *** |
| 10uM BMS202/$\gamma\delta$ T cells vs. ND/$\gamma\delta$ T cells | -24.01 | -39.22 to -8.789 | Yes | ** |
| 10uM BMS202/$\gamma\delta$ T cells vs. ND/2.5uM BMS202/$\gamma\delta$ T cells | -9.482 | -24.70 to 5.736 | No | ns |
| 10uM BMS202/$\gamma\delta$ T cells vs. ND/5uM BMS202/$\gamma\delta$ T cells | -3.397 | -18.61 to 11.82 | No | ns |
| 10uM BMS202/$\gamma\delta$ T cells vs. ND/10uM BMS 202/$\gamma\delta$ T cells | 13.16 | -2.061 to 28.37 | No | ns |
| ND/$\gamma\delta$ T cells vs. ND/2.5uM BMS202/$\gamma\delta$ T cells | 14.53 | -0.6921 to 29.74 | No | ns |
| ND/$\gamma\delta$ T cells vs. ND/5uM BMS 202/$\gamma\delta$ T cells | 20.61 | 5.393 to 35.83 | Yes | ** |
| ND/$\gamma\delta$ T cells vs. ND/10uM BMS202/$\gamma\delta$ T cells | 37.16 | 21.95 to 52.38 | Yes | *** |
| ND/2.5uM BMS202/$\gamma\delta$ T cells vs. ND/5uM BMS202/$\gamma\delta$ T cells | 6.085 | -9.133 to 21.30 | No | ns |
| ND/2.5uM BMS202/$\gamma\delta$ T cells vs. ND/10uM BMS202/$\gamma\delta$ T cells | 22.64 | 7.421 to 37.86 | Yes | ** |

## Notes

### Competing Interest Statement

The authors have declared no competing interest.

